# Genome-wide analysis of over 106,000 individuals identifies 9 neuroticism-associated loci

**DOI:** 10.1101/032417

**Authors:** Daniel J. Smith, Valentina Escott-Price, Gail Davies, Mark E.S. Bailey, Lucía Colodro-Conde, Joey Ward, Alexey Vedernikov, Riccardo Marioni, Breda Cullen, Donald Lyall, Saskia P. Hagenaars, David C.M. Liewald, Michelle Luciano, Catharine R. Gale, Stuart J. Ritchie, Caroline Hayward, Barbara Nicholl, Brendan Bulik-Sullivan, Mark Adams, Baptiste Couvy-Duchesne, Nicholas Graham, Daniel Mackay, Jonathan Evans, Blair H. Smith, David J. Porteous, Sarah Medland, Nick G. Martin, Peter Holmans, Andrew M. McIntosh, Jill P. Pell, Ian J. Deary, Michael O’Donovan

**Affiliations:** Institute of Health and Wellbeing, University of Glasgow, Glasgow, UK.; MRC Centre for Neuropsychiatric Genetics and Genomics, Cardiff University, Cardiff, UK.; Centre for Cognitive Ageing and Cognitive Epidemiology, Department of Psychology, University of Edinburgh, Edinburgh, UK.; School of Life Sciences, College of Medical, Veterinary and Life Sciences, University of Glasgow, Glasgow, UK.; Division of Psychiatry, University of Edinburgh, Edinburgh, UK.; QIMR Berghofer Medical Research Institute, Herston, Queensland, Australia.; Program in Medical and Population Genetics, Broad Institute of MIT and Harvard, Cambridge, Massachusetts, USA.; Analytical and Translational Genetics Unit, Department of Medicine, Massachusetts General Hospital and Harvard Medical School, Boston, Massachusetts, USA.; Stanley Center for Psychiatric Research, Broad Institute of MIT and Harvard, Cambridge, Massachusetts, USA.; MRC Lifecourse Epidemiology Unit, University of Southampton, Southampton General Hospital, Southampton, UK.; Medical Research Council Human Genetics Unit, Institute of Genetics and Molecular Medicine, University of Edinburgh, Edinburgh, UK.; Division of Population Health Sciences, University of Dundee, Dundee, UK.; Generation Scotland, Centre for Genomic and Experimental Medicine, Institute of Genetics and Molecular Medicine, University of Edinburgh, Edinburgh EH4 2XU, UK.; Medical Genetics Section, Centre for Genomic and Experimental Medicine, Institute of Genetics and Molecular Medicine, University of Edinburgh, Edinburgh EH4 2XU, UK.

## Abstract

Neuroticism is a personality trait of fundamental importance for psychological wellbeing and public health. It is strongly associated with major depressive disorder (MDD) and several other psychiatric conditions. Although neuroticism is heritable, attempts to identify the alleles involved in previous studies have been limited by relatively small sample sizes and heterogeneity in the measurement of neuroticism. Here we report a genome-wide association study of neuroticism in 91,370 participants of the UK Biobank cohort and a combined meta-analysis which includes a further 6,659 participants from the Generation Scotland Scottish Family Health Study (GS:SFHS) and 8,687 participants from a QIMR Berghofer Medical Research Institute (QIMR) cohort. All participants were assessed using the same neuroticism instrument, the Eysenck Personality Questionnaire-Revised (EPQ-R-S) Short Form’s Neuroticism scale. We found a SNP-based heritability estimate for neuroticism of approximately 15% (SE = 0.7%). Meta-analysis identified 9 novel loci associated with neuroticism. The strongest evidence for association was at a locus on chromosome 8 (p = 1.5x10^-15^) spanning 4 Mb and containing at least 36 genes. Other associated loci included interesting candidate genes on chromosome 1 (*GRIK3*, glutamate receptor ionotropic kainate 3), chromosome 4 (*KLHL2*, Kelch-like protein 2), chromosome 17 (*CRHR1*, corticotropin-releasing hormone receptor 1 and *MAPT*, microtubule-associated protein Tau), and on chromosome 18 (*CELF4*, CUGBP elav-like family member 4). We found no evidence for genetic differences in the common allelic architecture of neuroticism by sex. By comparing our findings with those of the Psychiatric Genetics Consortia, we identified a strong genetic correlation between neuroticism and MDD (0.64) and a less strong but significant genetic correlation with schizophrenia (0.22), although not with bipolar disorder. Polygenic risk scores derived from the primary UK Biobank sample captured about 1% of the variance in neuroticism in independent samples. Overall, our findings confirm a polygenic basis for neuroticism and substantial shared genetic architecture between neuroticism and MDD. The identification of 9 new neuroticism-associated loci will drive forward future work on the neurobiology of neuroticism and related phenotypes.

## Introduction

Neuroticism is a dimension of personality that has been studied for about 100 years, is present in most personality trait theories and questionnaires, and is found in the lexicons of most human cultures (1). Individual differences in neuroticism are highly stable across the life course (2). Higher neuroticism is associated with considerable public health and economic costs (3), premature mortality (4), and a range of negative emotional states and psychiatric disorders, including major depressive disorder (MDD), anxiety disorders, substance misuse disorders, personality disorders and schizophrenia (5-9). Thus, the study of neuroticism is not only important for understanding an important dimension of personality but may also illuminate the aetiology of a range of psychiatric disorders (10, 11).

H.J. Eysenck suggested a biological basis for neuroticism over 50 years ago (12). Although the biological underpinnings of personality traits are not understood, genetic factors are clearly involved. Twin studies suggest that about 40% of the trait variance for neuroticism is heritable (13-18), of which between 15-37% is explained by variation in common single nucleotide polymorphisms (SNPs) (18, 19) and is potentially detectable using the genome-wide association study (GWAS) paradigm. The clear links between neuroticism, psychopathology and other adverse health outcomes – and the implications for global health that would result from a better understanding of its mechanisms (20) – provide a strong rationale for large-scale GWAS to identify its genetic architecture (and genetic aetiology).

To date, individual GWAS of neuroticism have been limited by modest sample sizes and have delivered equivocal findings. Large meta-analyses of GWAS have also delivered modest findings. The recent Genetics of Personality Consortium (GPC) meta-analysis of neuroticism, which included 73,447 individuals from 29 discovery cohorts plus a replication cohort, identified only one genome-wide significant associated locus, at *MAGI1* on chromosome 3 (p=2.38×10^-8^) (19). Within two of the cohorts in this GPC study, common genetic variants explained approximately 15% of the variance in neuroticism (19).

In our study, seeking additional associated loci, we firstly used data from the UK Biobank cohort (21) to conduct a GWAS of neuroticism. Based on 91,370 participants from the UK, this is the largest single GWAS sample of neuroticism to date and the most homogeneous in terms of ascertainment strategy and assessment methodology. We then sought to extend these findings by conducting a meta-analysis which included the UK Biobank cohort, the Generation Scotland Scottish Family Health Study (GS:SFHS) cohort (22) and the QIMR Berghofer Medical Research Institute Study in Adults (QIMR) cohort (13-15). Additionally, we evaluated the genetic relationship between neuroticism and three major psychiatric phenotypes for which there are large, publically-accessible GWAS datasets: major depressive disorder (MDD); schizophrenia; and bipolar disorder (BD). Finally, we have compared our findings with those from the GPC meta-analytic GWAS of neuroticism (19), as well as the CONVERGE consortium for MDD (23).

## Materials and methods

### Sample

UK Biobank is a large prospective cohort of more than 502,000 residents of the United Kingdom, aged between 40 and 69 years (21). The aim of UK Biobank aim is to study the genetic, environmental, medication and lifestyle factors that cause or prevent disease in middle and older age. Recruitment occurred over a four-year period, from 2006 to 2010. Baseline assessments included social, cognitive, personality (the trait of neuroticism), lifestyle, and physical health measures. For the present study, we used the first genetic data release (June 2015) based on approximately one third of UK Biobank participants. Aiming to maximise homogeneity, we restricted the sample to those who reported being of white United Kingdom (UK) ancestry and for whom neuroticism phenotype data were available (n=91,370).

We also made use of data provided by investigators from the GS:SFHS (22) and QIMR cohorts (13-15) to conduct a meta-analysis based on samples for which we could readily access individual genotypes and which were assessed using the same measure of neuroticism. The GS:SFHS sample comprised 7,196 individuals and the QIMR sample comprised 8,687 individuals. Individuals (n = 537) who had participated in both UK Biobank and GS:SFHS were removed from the GS:SFHS sample based on relatedness checking using the genetic data.

Note that we were unable to incorporate the published data from the GPC as the neuroticism measure used in that study was derived from an item response theory (IRT) analysis (prohibiting inverse variance-weighted meta-analysis due to the differences in variance and heterogeneity of the measure). In addition, there was no information on the sample size for each SNP (prohibiting sample size-weighted meta-analysis) and the majority of participants in the QIMR cohort were included within the GPC meta-analysis.

This study obtained informed consent from all participants and was conducted under generic approval from the NHS National Research Ethics Service (approval letter dated 17th June 2011, Ref 11/NW/0382) and under UK Biobank approvals for application 6553 “Genome-wide association studies of mental health” (PI Daniel Smith) and 4844 “Stratifying Resilience and Depression Longitudinally” (PI Andrew McIntosh).

### Neuroticism phenotype

Neuroticism was assessed in all three cohorts (UK Biobank, GS:SFHS and QIMR) using the 12 items of the neuroticism scale from the Eysenck Personality Questionnaire-Revised Short Form (EPQ-R-S) (24) (supplementary table S1). Respondents answered ‘yes’ (score 1) or ‘no’ (score zero) to each of the questions, giving a total neuroticism score for each respondent of between 0-12. This short scale has a reliability of more than 0.8 (24) and high concurrent validity; for example, in a sample of 207 older people EPQ-R-S scores correlated 0.85 with the neuroticism score from the NEO-Five Factor Inventory, the scale most widely used internationally (25, 26).

### Genotyping and imputation

In June 2015 UK Biobank released the first set of genotype data for 152,729 UK Biobank participants. Approximately 67% of this sample was genotyped using the Affymetrix UK Biobank Axiom^®^ array and the remaining 33% were genotyped using the Affymetrix UK BiLEVE Axiom array. These arrays have over 95% content in common. Only autosomal data were available under the current data release. Data were pre-imputed by UK Biobank as fully described in the UK Biobank interim release documentation (27). Briefly, after removing genotyped single nucleotide polymorphisms (SNPs) that were outliers, or were multi-allelic or of low frequency (minor allele frequency, MAF < 1%), phasing was performed using a modified version of SHAPEIT2 and imputation was carried out using IMPUTE2 algorithms, as implemented in a C++ platform for computational efficiency (28, 29). Imputation was based upon a merged reference panel of 87,696,888 bi-allelic variants on 12,570 haplotypes constituted from the 1000 Genomes Phase 3 and UK10K haplotype panels (30). Variants with MAF < 0.001% were excluded from the imputed marker set. Stringent QC prior to release was applied by the Wellcome Trust Centre for Human Genetics (WTCHG), as described in UK Biobank documentation (31).

### Statistical analysis

#### Quality control and association analyses

Prior to all analyses, further quality control measures were applied. Individuals were removed based on UK Biobank genomic analysis exclusions (Biobank Data Dictionary item #22010), relatedness (#22012: genetic relatedness factor; a random member of each pair of individuals with KING-estimated kinship co-efficient > 0.0442 was removed), gender mismatch (#22001: genetic sex), ancestry (#22006: ethnic grouping; principal component analysis identified probable Caucasians within those individuals that were self-identified as British and other individuals were removed from the analysis) and QC failure in the UK BiLEVE study (#22050: UK BiLEVE Affymetrix quality control for samples and #22051: UK BiLEVE genotype quality control for samples). A sample of 112,031 individuals remained for further analyses. Of these, 91,370 had neuroticism scores. Genotype data were further filtered by removal of SNPs with Hardy-Weinberg equilibrium p<10^-6^, with MAF<0.01, with info<0.4, and with data on <95% of the sample after excluding genotype calls made with less than 90% posterior probability, after which 8,268,322 variants were retained.

Association analysis was conducted using linear regression under a model of additive allelic effects with sex, age, array, and the first 8 principal components (Biobank Data Dictionary items #22009.01 to #22009.08) as covariates. Genetic principal components (PCs) were included to control for hidden population structure within the sample, and the first 8 PCs, out of 15 available in the Biobank, were selected after visual inspection of each pair of PCs, taking forward only those that resulted in multiple clusters of individuals after excluding individuals self-reporting as being of non-white British ancestry (Biobank Data Dictionary item #22006). The distribution of the neuroticism score was assessed for skewness and kurtosis (coefficients were 0.56 and -0.61, respectively) and found to be sufficiently ‘normal’ (both coefficients are between -1 and 1) to permit analysis using linear regression. GWAS of neuroticism were additionally performed separately for females (N = 47,196) and males (N = 44,174) using linear regression (as above), with age, array, and the first 8 principal components as covariates.

#### Heritability, polygenicity, and cross-sample genetic correlation

Univariate GCTA-GREML analyses were used to estimate the proportion of variance explained by all common SNPs for the neuroticism phenotype (32). We additionally applied Linkage Disequilibrium Score Regression (LDSR)(33) to the summary statistics to estimate SNP heritability (h^2^_SNP_) and to evaluate whether inflation in the test statistics is the result of polygenicity or of poor control of biases such as population stratification. Genetic correlations between neuroticism scores in the three cohorts (UK Biobank, QIMR and GS:SFHS) were tested, and genetic correlations between neuroticism, schizophrenia, bipolar disorder (BD), and major depressive disorder (MDD) were evaluated in the UK Biobank sample using LDSR (34), a process that corrects for potential sample overlap without relying on the availability of individual genotypes (33). For the psychiatric phenotypes, we used GWAS summary statistics provided by the Psychiatric Genomics Consortium (http://www.med.unc.edu/pgc/) (35-37).

#### Polygenic risk score analyses in the QIMR and GS:SFHS samples

In the QIMR sample (N = 8,687 individuals), Polygenic Risk Scores for neuroticism (PRS-N) based on the summary statistics from the UK Biobank GWAS were computed with PLINK 1.90 (version Sep 3rd 2015, https://www.cog-genomics.org/plink2/), for p value thresholds (PT) 0.01, 0.05, 0.1, 0.5, and 1; following the procedure described by Wray and colleagues (38). All subjects had GWAS data imputed to 1000G v.3. Only SNPs with a minor allele frequency ≥0.01 and imputation quality r^2^≥0.6 were used in the calculation of the PRS-N. Genotypes were LD pruned using clumping to obtain SNPs in approximate linkage equilibrium with an r^2^<0.1 within a 10,000bp window. Since QIMR participants were related, predictions were calculated using GCTA (Genome-wide Complex Trait Analysis, version 1.22)(39), using the following linear mixed model: EPQ-N = intercept + beta0^*^ covariates + beta2 ^*^ g + e with g^~^N(0, GRM), where: EPQ is neuroticism measured by EPQ (standardised sum score); covariates are age, sex, imputation chip, ten genetic principal components and the standardised PRS (PT 0.01, 0.05, 0.1, 0.5, or 1); e is error; and GRM is genetic relationship matrix. P-values were calculated using the t-statistic on the basis of the Beta and SE from the GCTA output. Variance explained by the PRS was calculated using: var(x)^*^b^2/var(y), where x is the PRS, b is the estimate of the fixed effect from GCTA and y is the phenotype.

In the GS:SFHS sample, PRS-N based on the UK Biobank neuroticism GWAS results were created using PRSice from observed genotypes in 7,196 individuals (22, 40). SNPs with a minor allele frequency <0.01 were removed prior to creating PRS-N. Genotypes were LD pruned using clumping to obtain SNPs in linkage equilibrium with an r^2^<0.25 within a 200kb window. As above, five PRS-N were created containing SNPs according to the significance of their association with the phenotype, with PTs of 0.01, 0.05, 0.1, 0.5, and 1 (all SNPs). Linear regression models were used to examine the associations between the PRS-N and neuroticism score in GS, adjusting for age at measurement, sex and the first 10 genetic principal components to adjust for population stratification. The False Discovery Rate method was used to correct for multiple testing across the PRS-N at all five thresholds (41).

#### Meta-analysis

Inverse variance-weighted meta-analysis of UK Biobank, GS:SFHS and QIMR results was performed, restricted to variants present in all 3 samples, using the METAL package (http://www.sph.umich.edu/csg/abecasis/Metal). Data were available across all 3 studies for 7,207,648 of the original 8,268,322 variants from the UK Biobank analysis. The total sample size included in the meta-analysis was N = 106,716 (UK Biobank N = 91,370; GS:SFHS N = 6,659; QIMR N = 8,687).

## Results

### Neuroticism phenotype within UK Biobank and sociodemographic characteristics

Sociodemographic details of the 91,370 UK Biobank participants used in this analysis, as well as the full UK Biobank sample, are provided in table 1 and the distributions of neuroticism scores for males and females in our sample are provided in figure 1. The proportion of the UK Biobank neuroticism GWAS sample holding a degree was 31.4%, and the mean age of leaving full-time education for those without a degree was 16.5 years. Those in the full UK Biobank sample who responded to the neuroticism questions tended to be better educated than those who did not (33.4% had an undergraduate degree versus 27.7% in non-responders). As expected (42), mean neuroticism scores were lower for men than for women (men mean EPQ-R-S = 3.58, SD = 3.19; women mean EPQ-R-S = 4.58, SD = 3.26; p = 0.001). Principal component analysis of the 12 EPQ-R-S items showed that all items loaded highly on a single component, and the internal consistency (Cronbach alpha) coefficient was 0.84 (supplementary table S2). Analysis of the entire UK Biobank sample (N with data = 401,695) gave very similar results (supplementary table S2), suggesting the subsample analysed here is representative of the whole UK Biobank cohort.

**Table 1.**
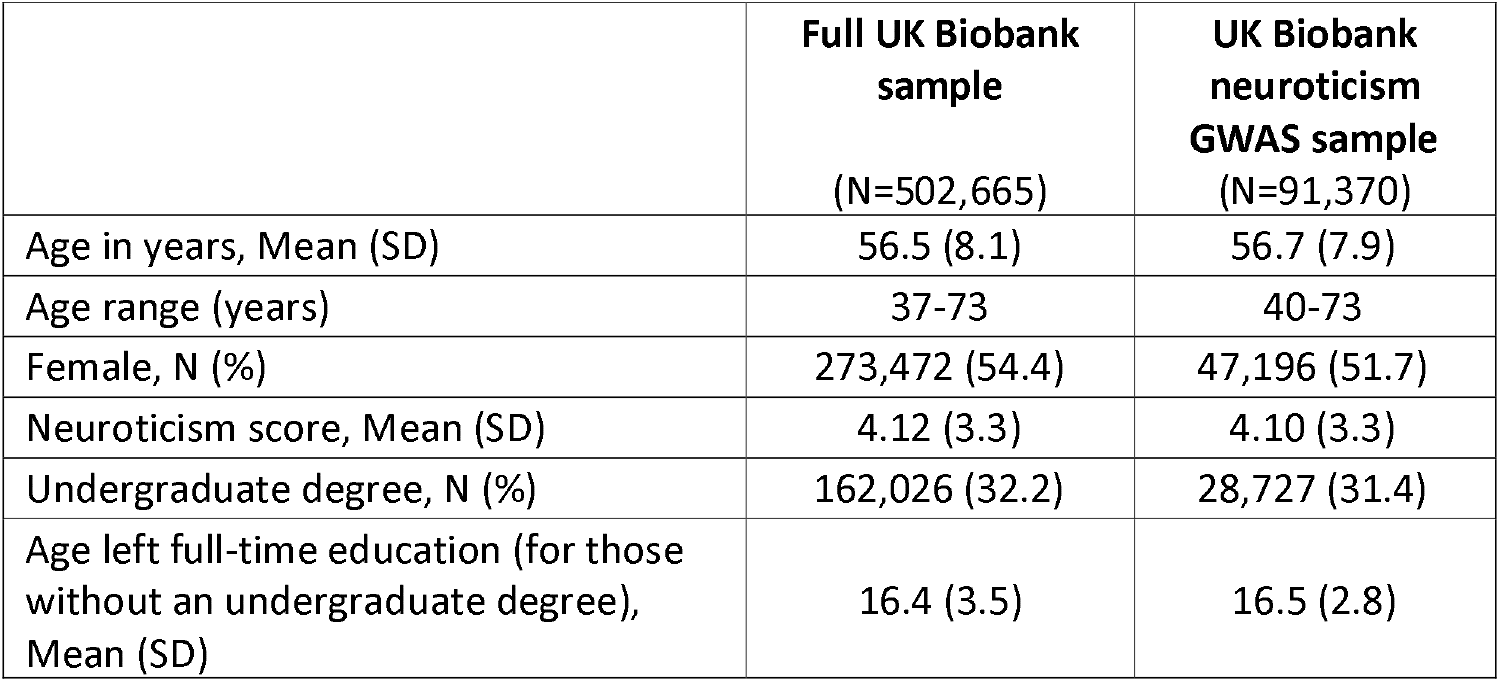
Sociodemographic characteristics in UK Biobank.

|  | **Full UK Biobank**<br>**sample**<br>(N=502,665) | **UK Biobank**<br>**neuroticism**<br>**GWAS sample**<br>(N=91,370) |
| --- | --- | --- |
| Age in years, Mean (SD) | 56.5 (8.1) | 56.7 (7.9) |
| Age range (years) | 37-73 | 40-73 |
| Female, N (%) | 273,472 (54.4) | 47,196 (51.7) |
| Neuroticism score, Mean (SD) | 4.12 (3.3) | 4.10 (3.3) |
| Undergraduate degree, N (%) | 162,026 (32.2) | 28,727 (31.4) |
| Age left full-time education (for those<br>without an undergraduate degree),<br>Mean (SD) | 16.4 (3.5) | 16.5 (2.8) |

**Figure 1.**
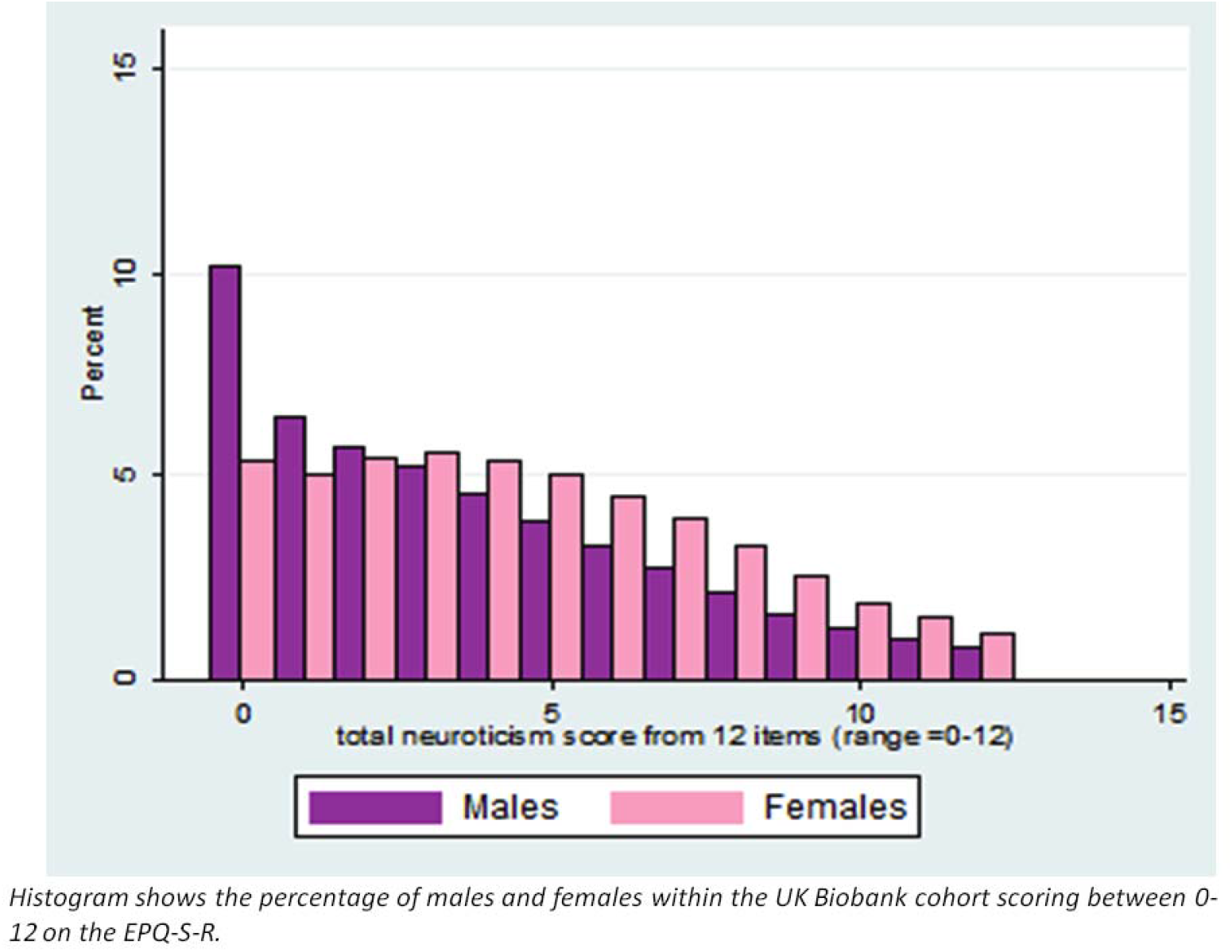
Distribution of neuroticism scores in UK Biobank sample (n=91,370).

### Genome-wide association results in UK Biobank

Genome-wide association results from the UK Biobank cohort are summarized in supplementary materials: supplementary figure S1 (QQ plot); supplementary figure S2 (Manhattan plot); and supplementary table S3 (genome-wide significant loci associated with neuroticism).

Overall, the GWAS data showed modest deviation in the test statistics compared with the null (λ_GC_ = 1.152); this was negligible in the context of sample size (λ_GC_1000 = 1.003) (figure S1). LDSR (33) suggested that deviation from the null was due to a polygenic architecture in which h^2^_SNP_ accounted for about 14% of the population variance in neuroticism (liability scale h^2^_SNP_ = 0.136 (SE 0.0153)), rather than inflation due to unconstrained population structure (LD regression intercept = 0.982 (SE 0.014)). Estimates of heritability using GCTA were similar to those using LD score regression (h^2^ = 0.156, SE = 0.0074).

We observed a total of 8 independent loci exhibiting genome-wide significant associations with neuroticism (figure S2, supplementary table S3) with the strongest evidence for association coming from a locus on chromosome 8 (p = 1.02x10^-15^) at which there is an extensive LD block spanning 4 Mb (attributable to an inversion polymorphism which has suppressed recombination) containing at least 36 genes. Similar findings to those from the UK Biobank dataset in a GWAS primarily assessing the genetics of wellbeing have also recently been posted in a non-peer reviewed format (43).

### Stratification by sex in UK Biobank

Neuroticism scores are in general higher in women than in men and it has been postulated that neuroticism may play a stronger etiologic role in MDD in women than in men (44, 45), potentially explaining the greater prevalence of depressive and anxiety disorders in women (46). This suggests the possibility of sex-related genetic heterogeneity. We therefore conducted secondary analyses looking for sex-specific neuroticism loci in women (N = 47,196) and men (N = 44,174) respectively. To minimize heterogeneity, this analysis was restricted to the UK Biobank samples. SNP heritability (measured by LDSR) for each sex was comparable (female h^2^_SNP_ = 0.149 (SE = 0.0169); male h^2^_SNP_ = 0.135 (SE = 0.0237)), and was highly correlated between the sexes (genetic correlation = 0.911 (SE = 0.07); p = 1.07x10^-38^) at a level that was not significantly different from 1 (p=0.21). In both sexes separately, the chromosome 8 locus was associated at genome-wide significance but no other single locus attained significance. Overall, we found no evidence for genetic differences in the common allelic architecture of neuroticism by sex.

### Meta-analysis of UK Biobank, GS:SFHS and QIMR samples

In the combined dataset, we obtained genome wide significance for 9 independent loci: on chromosome 1 (two loci), chromosome 3, chromosome 4, chromosome 8, chromosome 9 (two loci), chromosome 17 and chromosome 18 (figure 2, table 2).

**Figure 2.**
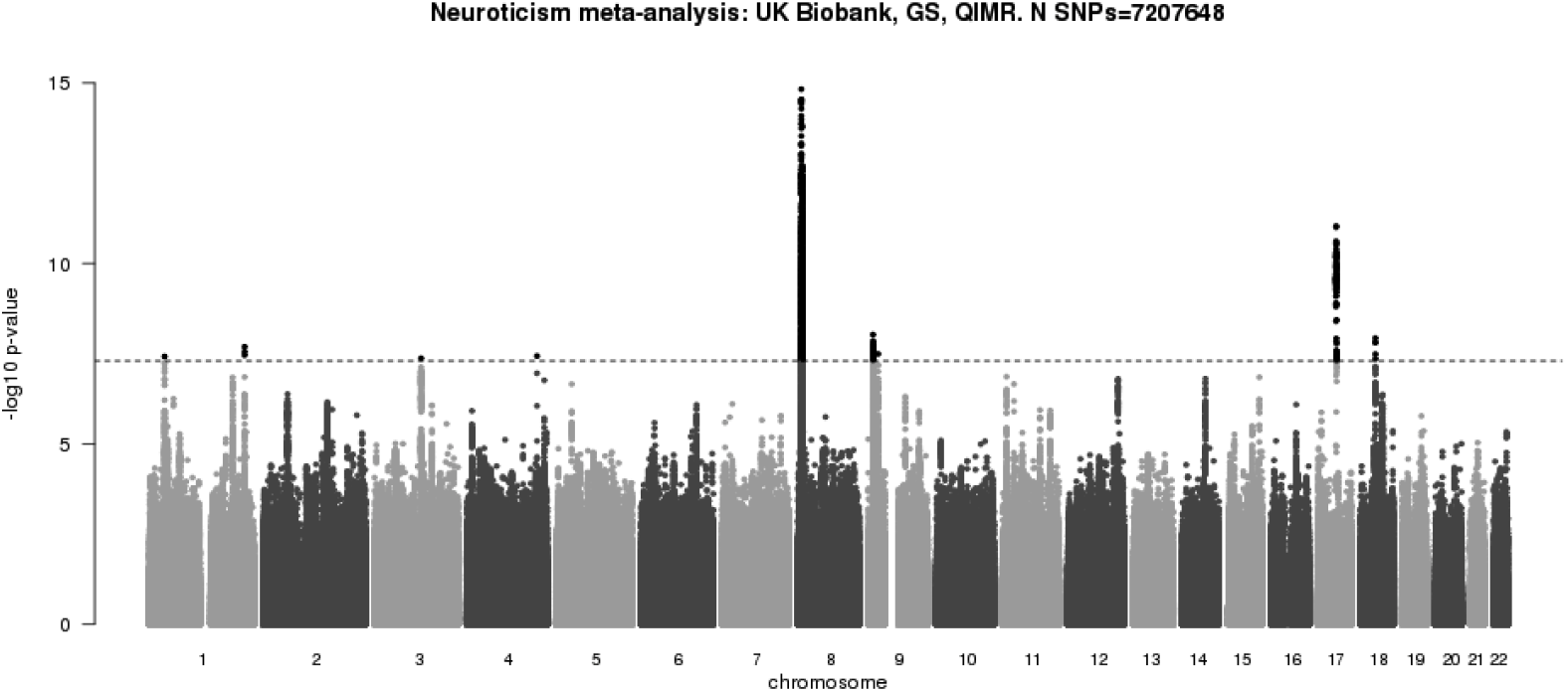
Manhattan plot of meta-analysis of GWAS from UK Biobank, GS:SFHS and QIMR samples (combined N = 106,716).

**Table 2.**
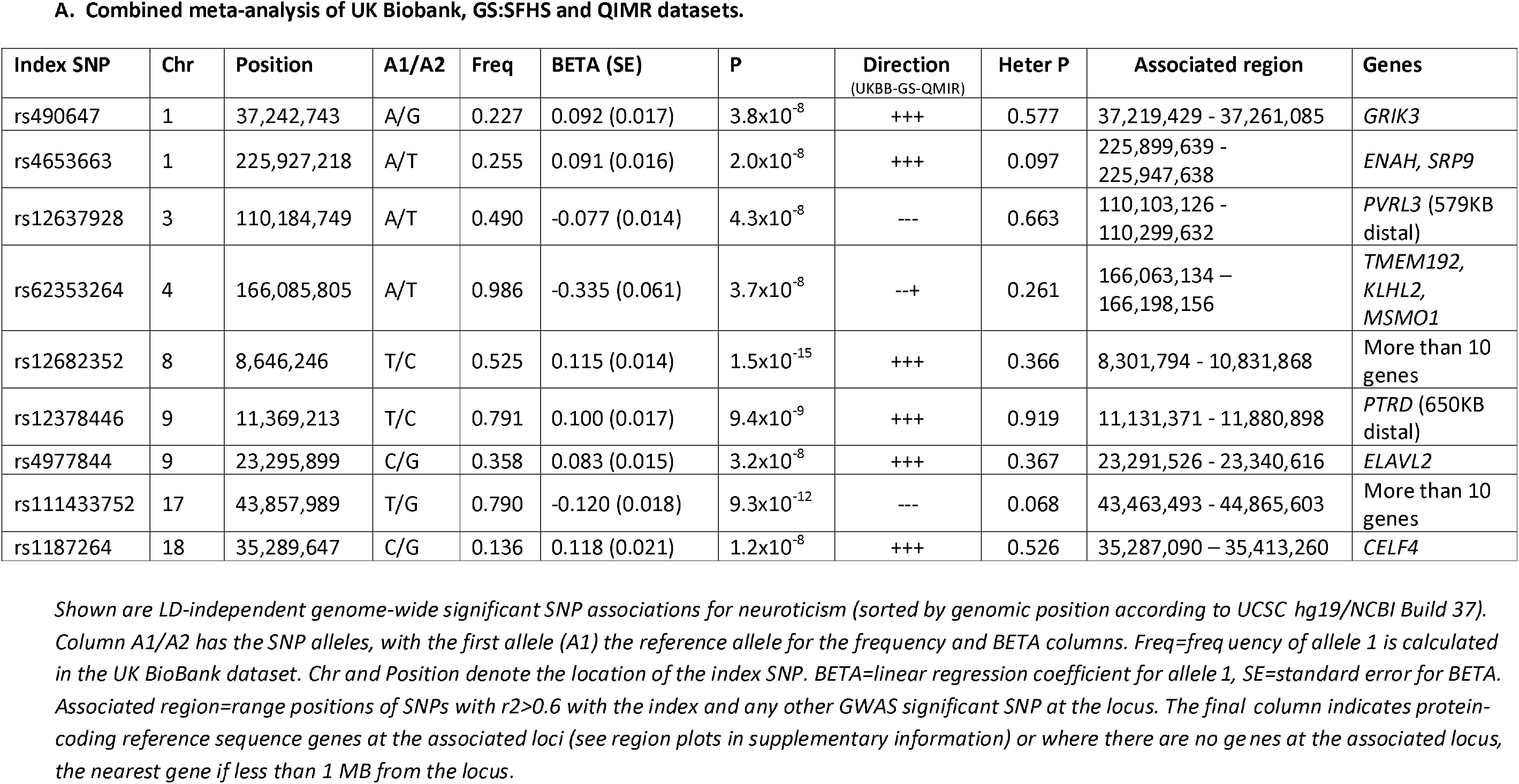
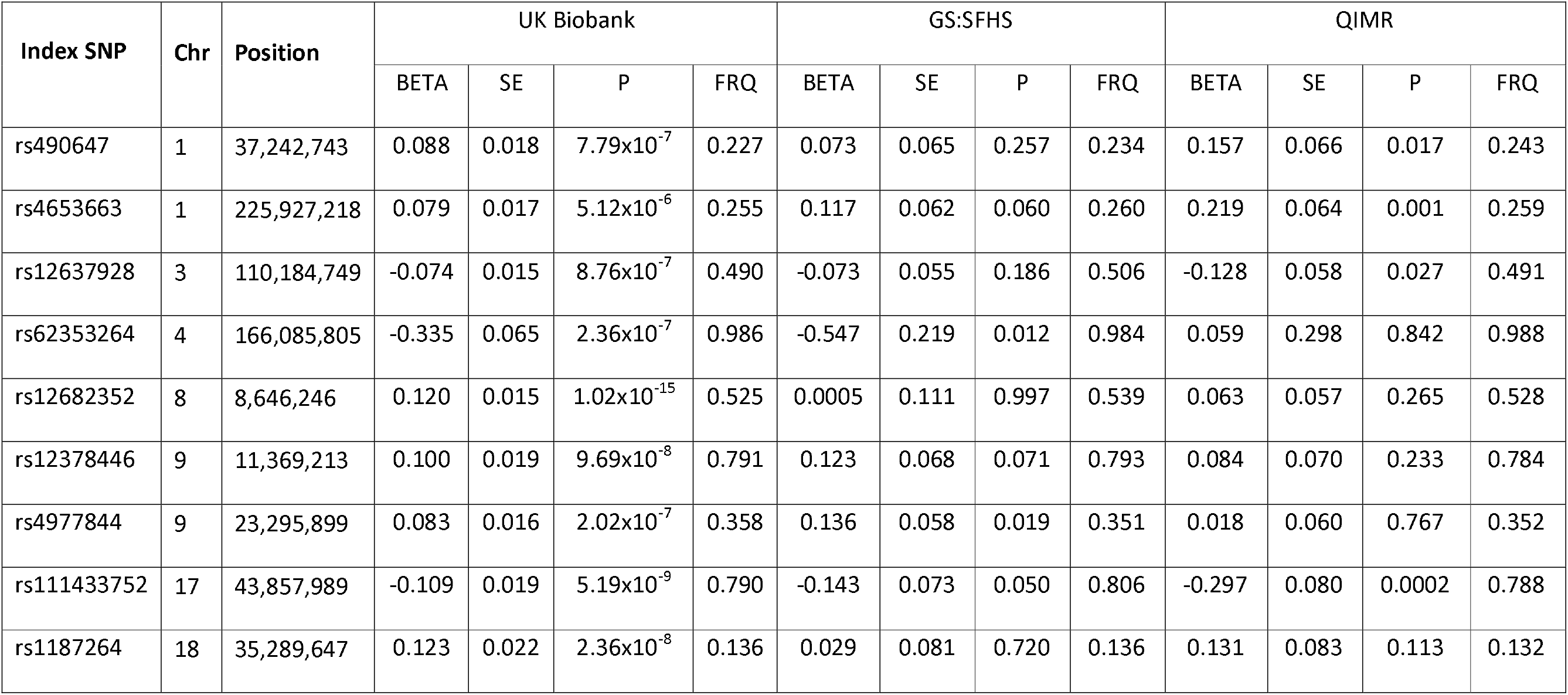
Genome-wide significant index SNPs.

Full details are provided in table 2 and the associated regions are depicted graphically as region plots in supplementary figure S3 (S3a-S3i). Candidate genes of particular note mapping to the associated loci include: the glutamatergic kainate receptor *GRIK3* (figure S3a) (47, 48); *CELF4*, which regulates excitatory neurotransmission (figure S3i) (49); and *CRHR1*, encoding corticotropin-releasing hormone receptor 1 (figure s3h), a protein that is central to the stress response (50). Associated loci are considered in greater detail within the discussion.

### Genetic correlation of neuroticism with MDD, schizophrenia and bipolar disorder

LDSR showed strong genetic correlation between neuroticism and MDD (genetic correlation= 0.64, SE = 0.071, p = 3.31x10^-19^) and a smaller, but significant, correlation between neuroticism and schizophrenia (genetic correlation = 0.22, SE = 0.05, p = 1.96x10^-05^) (table 3). We found no significant overlap between neuroticism and bipolar disorder (genetic correlation = 0. 07, SE = 0.05, p = 0.15). Similar results based solely on the UK Biobank dataset have been reported recently in a non-peer reviewed format (51).

**Table 3.** Genetic correlations between neuroticism and MDD, schizophrenia and bipolar disorder.

|  | N cases | N controls | Genetic Correlation | SE Genetic correlation | Significance (p-value) |
| --- | --- | --- | --- | --- | --- |
| MDD | 9240 | 9519 | 0.64 | 0.07 | $3.31 \times 10^{-19}$ |
| Bipolar disorder | 7481 | 9250 | 0.07 | 0.05 | 0.1505 |
| Schizophrenia | 34241 | 45604 | 0.22 | 0.05 | $1.96 \times 10^{-5}$ |

Columns “N cases” and “N controls” show the numbers of cases and controls in the corresponding PGC2 genome-wide association studies (https://www.med.unc.edu/pgc/downloads). Columns 4,5,6 present genetic correlation estimates, their standard errors and significance, respectively, calculated with LD Score regression tool (https://github.com/bulik/ldsc).

### Genetic correlations for neuroticism between UK Biobank, GS:SFHS and QIMR samples

The LDSR-calculated genetic correlation for neuroticism between the three samples was strong: between UK Biobank and GS:SFHS the genetic correlation was 0.91 (SE = 0.15, p = 4.04x10^-09^); between UK Biobank and QIMR the genetic correlation was 0.74 (SE = 0.14, p = 2.49x10^-07^), and between GS:SFHS and QIMR the genetic correlation was 1.16 (SE = 0.35, p = 0.0009). Note that the true maximum for a genetic correlation is bounded by 1. That the LD score estimate is greater than this reflects the imprecision in the estimate as indicated by the large SE, in the context of which we interpret this as evidence for high but imprecisely estimated genetic correlation between the two samples.

### Polygenic risk score (PRS) analysis for neuroticism in GS:SFHS and QIMR samples

Table 4 shows the results of PRS analysis (based on the UK Biobank-only GWAS) within the GS:SFHS and QIMR samples. At all thresholds tested, PRS-N predicted neuroticism, although the amount of variance explained was small (at around 1%).

**Table 4.** Associations between the polygenic risk scores (PRS) for Neuroticism based on the UK Biobank Neuroticism GWAS summary results, and Neuroticism in GS:SFHS and QIMR samples, controlling for age, sex, and ten genetic principal components for population structure.

| **GS:SFHS**<br>**sample**<br>**N = 7,196** |  |  |  |  |  |
| --- | --- | --- | --- | --- | --- |
| **Threshold** | **Beta** | **SE** | **Percentage**<br>**variance**<br>**explained** | **P value** | **Number of SNPs** |
| PRS<0.01 | 0.107 | 0.016 | 0.59 | $4.58 \times 10^{-11}$ | 4531 |
| PRS<0.05 | 0.123 | 0.014 | 1.00 | $5.27 \times 10^{-19}$ | 15533 |
| PRS<0.1 | 0.131 | 0.013 | 1.30 | $3.23 \times 10^{-23}$ | 27216 |
| PRS<0.5 | 0.132 | 0.012 | 1.48 | $3.45 \times 10^{-26}$ | 95552 |
| PRS<1 | 0.131 | 0.012 | 1.46 | $6.93 \times 10^{-26}$ | 146088 |
| **QIMR**<br>**Sample**<br>**N = 8,687** |  |  |  |  |  |
| **Threshold** | **Beta** | **SE** | **Percentage**<br>**variance**<br>**explained** | **P value** | **Number of SNPs** |
| PRS<0.01 | 0.070 | 0.012 | 0.49 | $8.5 \times 10^{-9}$ | 12,146 |
| PRS<0.05 | 0.081 | 0.012 | 0.66 | $5.3 \times 10^{-12}$ | 41,006 |
| PRS<0.1 | 0.086 | 0.012 | 0.74 | $1.5 \times 10^{-13}$ | 68,979 |
| PRS<0.5 | 0.086 | 0.012 | 0.75 | $7.7 \times 10^{-14}$ | 204,632 |
| PRS<1 | 0.088 | 0.011 | 0.77 | $3.2 \times 10^{-14}$ | 280,716 |

### Comparison with findings from GPC meta-analysis

In contrast to the finding of the GPC meta-analysis, we did not identify a genome-wide significant hit close to *MAGI1* within 3p14 (19). However, within the UK Biobank sample, the same allele at the associated SNP from that study (rs35855737) did show a trend for association (Beta = 0.035, SE = 0.02, p = 0.07; 2-tailed) in the expected direction.

### Comparison with findings from the CONVERGE consortium study of MDD

The recently-published CONVERGE consortium study of Chinese women with recurrent and melancholic MDD identified two loci contributing to risk of MDD on chromosome 10; one near the *SIRT1* gene (rs12415800; P = 2.53 × 10^-10^), the other in an intron of the *LHPP* gene (rs35936514, P = 6.45 × 10^-12^) (23). Neither of these index SNPs were associated with neuroticism within the UK Biobank sample (for rs12415800 Beta = -0.107, SE = 0.066, p = 0.1036, freq A=0.013; and for rs35936514 Beta = 0.021, SE = 0.0378, p = 0.5832, freq T= 0.041).

## Discussion

The identification of 9 independent loci showing genome-wide significant associations with neuroticism within our combined meta-analysis represents a significant advance. In contrast, a recent meta-analysis of neuroticism conducted by the GPC (n = 73,447) identified only a single genome-wide significant locus (19).

There are several possible explanations for this difference. All three of the cohorts in our study used the same reliable and validated 12-item neuroticism assessment instrument (EPQ-R-S), whereas the GPC study assessed neuroticism scores collected using several different instruments across thirty cohorts. Although the GPC used item response theory (IRT) analysis to harmonise neuroticism scores (18), this is likely to be much less reliable than the use of a single consistent instrument. Further, the UK Biobank cohort is by far the largest sample ever studied for neuroticism genetics and all of the participants were of White British ethnicity, minimising population stratification and also addressing potential problems with cultural variation in the interpretation of neuroticism questionnaire items. Additionally, quality control steps in the UK Biobank sample were performed in a single centre in a consistent way.

The most significant associated locus on chromosome 8, which was independently associated at genome-wide significance for both men and women, spans a 4 Mb region of extended LD (the result of an inversion polymorphism) containing at least 36 genes (table 2 and supplementary figure S3e). The extended LD at this locus means that identifying the specific genes responsible for the association is likely to prove challenging. As an initial attempt to resolve the signal, we queried the index SNP (rs12682352) at the BRAINEAC (http://www.braineac.org/) brain eQTL resource. This identified *ERI1* as the only protein coding gene within the locus whose expression was associated with the index SNP in brain, but only nominally so (p=0.019) and not at a level that would reliably point to this gene as likely explaining the association.

The locus on chromosome 17 (rs111433752 at 43.8 MB; supplementary figure S3h) similarly maps to an inversion polymorphism spanning multiple genes and therefore we cannot attribute the association to any particular gene. As with the locus on chromosome 8, inspection of eQTLs in the region in BRAINEAC did not help to resolve the signal. Nevertheless, this locus contains a notable candidate gene, *CRHR1*, encoding corticotropin-releasing hormone receptor 1. In the presence of corticotropin-releasing hormone (CRH), *CRHR1* triggers the downstream release of the stress response-regulating hormone cortisol. CRHR1 is therefore a key link in the hypothalamic-pituitary-adrenal (HPA) pathway which mediates the body’s response to stress and which is abnormal in severe depression (50). *CRHR1* per se has also been shown to be involved in anxiety-related behaviours in mice and has also been genetically associated with panic disorder in humans (52).

Another potential candidate gene within the extended region of genome-wide significant association at the chromosome 17 locus is *MAPT*, which encodes the microtubule-associated protein Tau. There is evidence that Tau is present in the postsynaptic compartment of many neurons (53) and *MAPT* knockout in mice leads to defects in hippocampal long-term depression (LTD) (54), as well as mild network-level alterations in brain function (55). The clearest candidate gene at one of the other loci, *CELF4* on chromosome 18 at approximately 35Mb, encodes an mRNA binding protein known to participate in a major switch in Tau protein isoform distribution after birth in the mammalian brain (56). CELF4 is expressed predominantly in glutamatergic neurones, and recent studies suggest it has a central role in regulating excitatory neurotransmission by modulating the stability and/or translation of a range of target mRNAs (49).

The finding of an association with a locus on chromosome 1 (rs490647), which includes the glutamatergic kainate receptor *GRIK3*, is of considerable interest given that abnormalities of the glutamate system are implicated in the pathophysiology of MDD (57-62). Further, a recent glutamate receptor gene expression study in a large cohort of post-mortem subjects, including some individuals with MDD who had completed suicide, found *GRIK3* to be the strongest predictor of suicide (48).

On chromosome 4, rs62353264 lies a short distance upstream of *KLHL2*, which encodes a BTB-Kelch-like protein. KLHL2 is an actin-binding protein and has also been reported to be part of a complex that ubiquitinates NPTXR, the neuronal pentraxin receptor (63), amongst other targets. Expression of KLHL2 has been reported to be enriched in brain, and it is localised to cytoplasm and processes of neurons and astrocytes, being found at sites of ruffles and other actin network-containing membrane outgrowths (64, 65). The associated region at this locus is short (approximately 150kb), and although several other genes lie within 500kb of the peak association at this locus, none is as promising a candidate as *KLHL2*.

The associated region in chromosome 9p23, at around 11.2-11.7Mb contains no protein-coding genes; the nearest gene on the telomeric side, with its 5’-end located about 650 kb from the associated region, is *PTPRD*. This gene encodes a receptor-type protein tyrosine phosphatase known to be expressed in brain and with an organising role at a variety of synapses (66), including those that play a role in synaptic plasticity. *PTPRD* is also known to harbour variation associated with restless legs syndrome (67). This is a credible candidate but particular caution is required given the distance between the associated locus and this gene.

In addition to identifying genome-wide significant loci, our study contributes further to understanding the genetic architecture of neuroticism and its relationship to other disorders. Our SNP-based heritability estimate for neuroticism was around 0.15, as estimated using GCTA, and only slightly lower using LDSR. This is consistent with the estimates reported by the GPC (19) in the two homogeneous subsets of the data they tested, and considerably greater than some earlier reports of approximately 6% (68, 69). Despite differences in the distribution of neuroticism by sex, SNP-based heritability was similar for both men and women and the genetic correlation between sexes was not significantly different from 1, suggesting a similar common variant architecture for both, and that differences in trait scores between the sexes are likely to result from structural variants, rare alleles and/or environmental exposures.

PRS analysis of neuroticism within the GS:SFHS and QIMR samples supported the expected highly polygenic architecture of neuroticism; despite the large discovery UK Biobank sample – but consistent with the modest number of GWS findings identified in this large sample – extremely weakly associated alleles at relaxed association thresholds (e.g., P_T_ up to at least 0.5) contributed to the variance captured by the signal.

Consistent with current practice, we regard the meta-analysis results as the primary outputs of this study. However, it is notable that while the results of the polygenic risk score analyses show that *en masse*, alleles that associate with neuroticism in UK Biobank tend to do the same in those with higher neuroticism within GS:SFHS and QIMR, this is not evident for the loci attaining genome-wide significance. Thus, of the 8 loci that were genome-wide significant in the UK Biobank dataset, only 5 remain significantly associated within the meta-analysis. With the exception of the locus on chromosome 17, none of these are strongly or consistently replicated across the GS:SFHS and QIMR samples, and the most significantly associated locus, that on chromosome 8, is not significant in either sample (supplementary table S4). The large standard errors for the estimates of effect sizes in GS:SFHS and QIMR are consistent with low power of these population samples to detect loci (with the effect sizes seen in complex traits), and with the fact that fully independent replication (or refutation) will require much larger samples.

By comparing the overall association analysis results in our study with those from the Psychiatric Genomics Consortia, we identified a strong genetic correlation between neuroticism and MDD (0.64), and a weaker but still significant genetic correlation with schizophrenia (0.22), although not with bipolar disorder. These findings are line with evidence suggesting that neuroticism and MDD –as well as, to a lesser extent, neuroticism and schizophrenia – share genetic risk factors in common (70). However, the present findings do not distinguish between a direct causal link between neuroticism and those other disorders (5, 7, 8, 71) versus pleiotropy, whereby a proportion of risk alleles that influence neuroticism also exert an effect on the clinical diagnoses. Nevertheless, our findings suggest neuroticism as a potentially fruitful measure for efforts such as the Research Domain Criteria (RDoC) initiative that seek to use fundamental and quantitative characteristics to investigate the etiology of psychiatric disorders across traditional nosological boundaries, in order to develop a more biologically-informed system of psychiatric classification (72).

Our findings are of interest in the context of the limited success to date of GWAS studies of MDD. A recent mega-analysis of genome-wide association studies for MDD (9,240 MDD cases and 9,519 controls in discovery phase, and 6,783 MDD cases and 50,695 controls in replication phase) failed to identify any genome-wide significant SNPs, suggesting that much larger samples are required to detect genetic effects for complex traits such as MDD (37). Given the high genetic correlation between neuroticism and MDD, combining the two datasets in a meta-analysis may be a plausible strategy to optimise the power of population samples in the search for a proportion of MDD loci, while noting that the two phenotypes are not perfectly genetically correlated. The MDD loci identified in a recent study of Chinese women with recurrent (N = 5,303) and melancholic (N = 4,509) MDD by the CONVERGE consortium (23) did not overlap with any of the loci reported here; given the apparent modest power to detect genome-wide significant loci in our sample, population differences between the studies and substantial differences between the phenotypes, the absence of overlap does not provide any evidence against the validity of the CONVERGE study finding. Given that neuroticism is a personality trait established as phenotypically and genetically strongly associated with MDD, the identification of several new genome-wide significant loci for neuroticism represents an important potential entry point into the biology of MDD.

### Conclusion

Overall, our findings confirm a polygenic basis for neuroticism and substantial shared genetic architecture between neuroticism and MDD, and to a lesser extent with schizophrenia, though not with bipolar disorder. The identification of 9 new loci associated with neuroticism represents a significant advance in this field and will drive future work on the neurobiology of a personality trait which has fundamental importance to human health and wellbeing.

## Acknowledgements

This research was conducted using the UK Biobank resource. UK Biobank was established by the Wellcome Trust, Medical Research Council, Department of Health, Scottish Government and Northwest Regional Development Agency. UK Biobank has also had funding from the Welsh Assembly Government and the British Heart Foundation. Data collection was funded by UK Biobank. Generation Scotland (GS:SFHS) receives core support from the Chief Scientist Office of the Scottish Government Health Directorates (CZD/16/6) and the Scottish Funding Council. We are grateful to all the families who took part, the general practitioners and the Scottish School of Primary Care for their help in recruiting them, and the whole Generation Scotland team, which includes interviewers, computer and laboratory technicians, clerical workers, research scientists, volunteers, managers, receptionists, healthcare assistants and nurses. DJS is supported by an Independent Investigator Award from the Brain and Behaviour Research Foundation (21930). AMM, IJD and MA are supported by Welcome Trust Strategic Award 104036/Z/14/Z. LCC is supported by a post-doctoral fellowship from the Fundación Séneca (Seneca Foundation, Regional Agency for Science and Technology, Murcia, Spain, 19151/PD/13). SEM is supported by a Future Fellowship from the Australian Research Council. The funders had no role in the design or analysis of this study, decision to publish, or preparation of the manuscript. We acknowledge support (QIMR study) from Grant W. Montgomery and Andrew C. Heath.

## Figures in supplementary material

**Figure S1.**
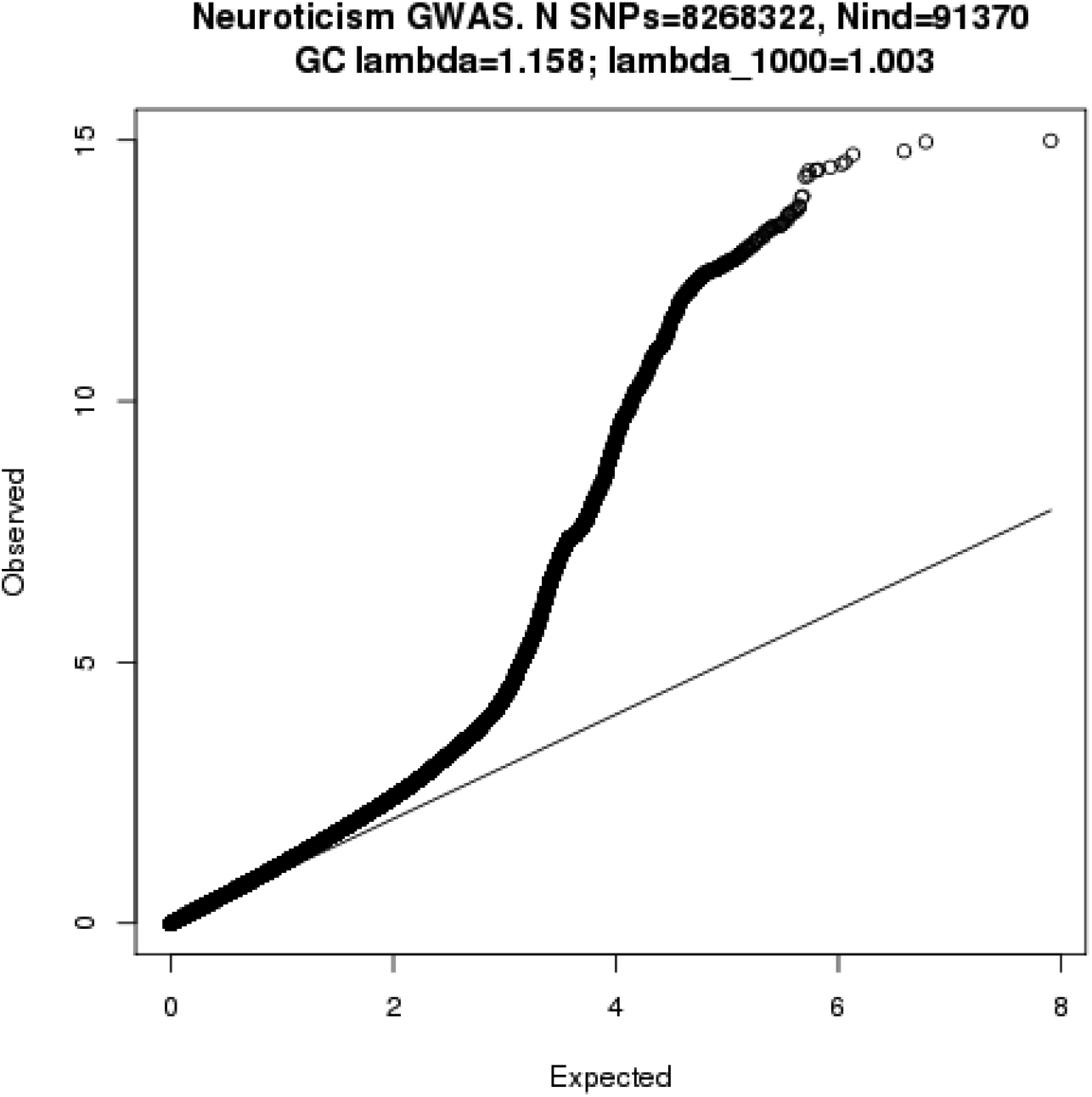
QQ plot for genome-wide association with neuroticism (n=91,370 UK Biobank participants only).

**Figure S2.**
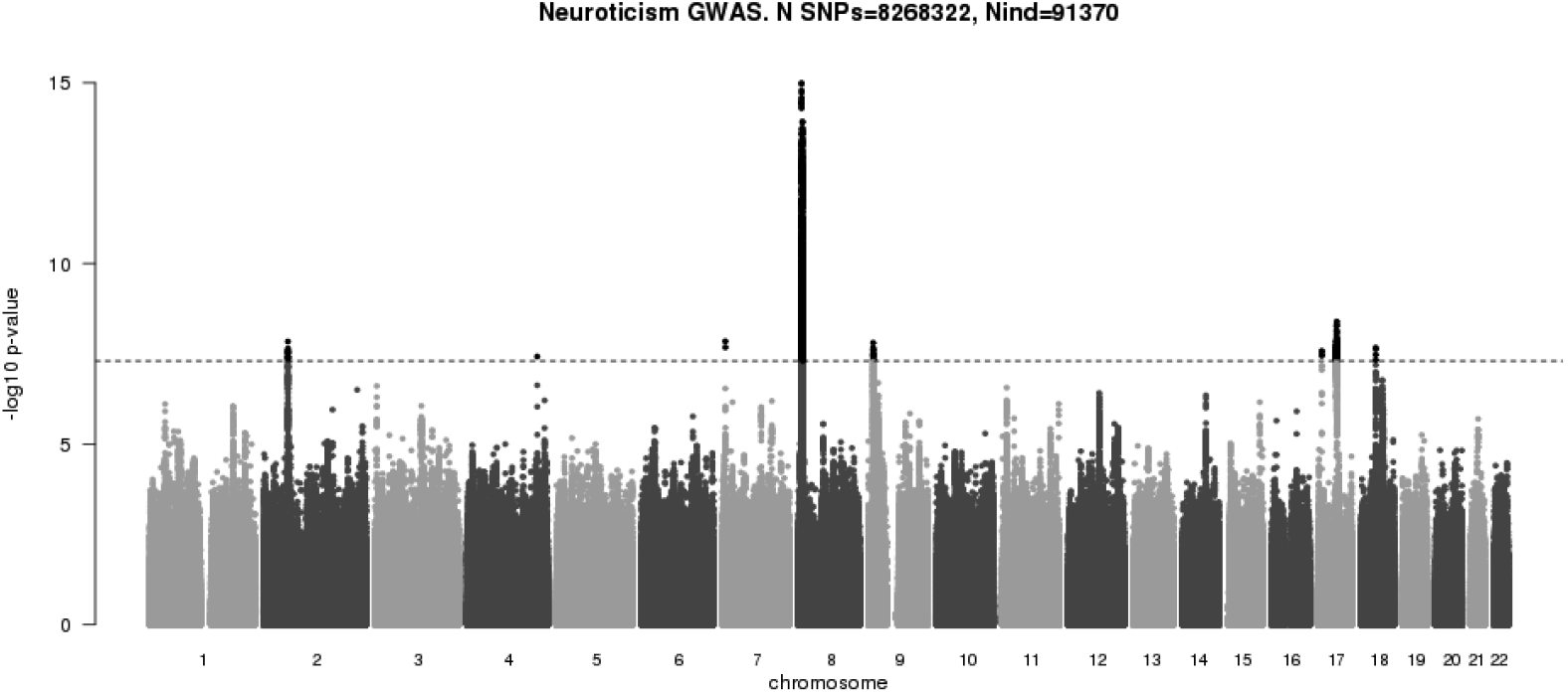
Manhattan plot (GWAS of N = 91,370 UK Biobank participants only).

**Figure S3.**
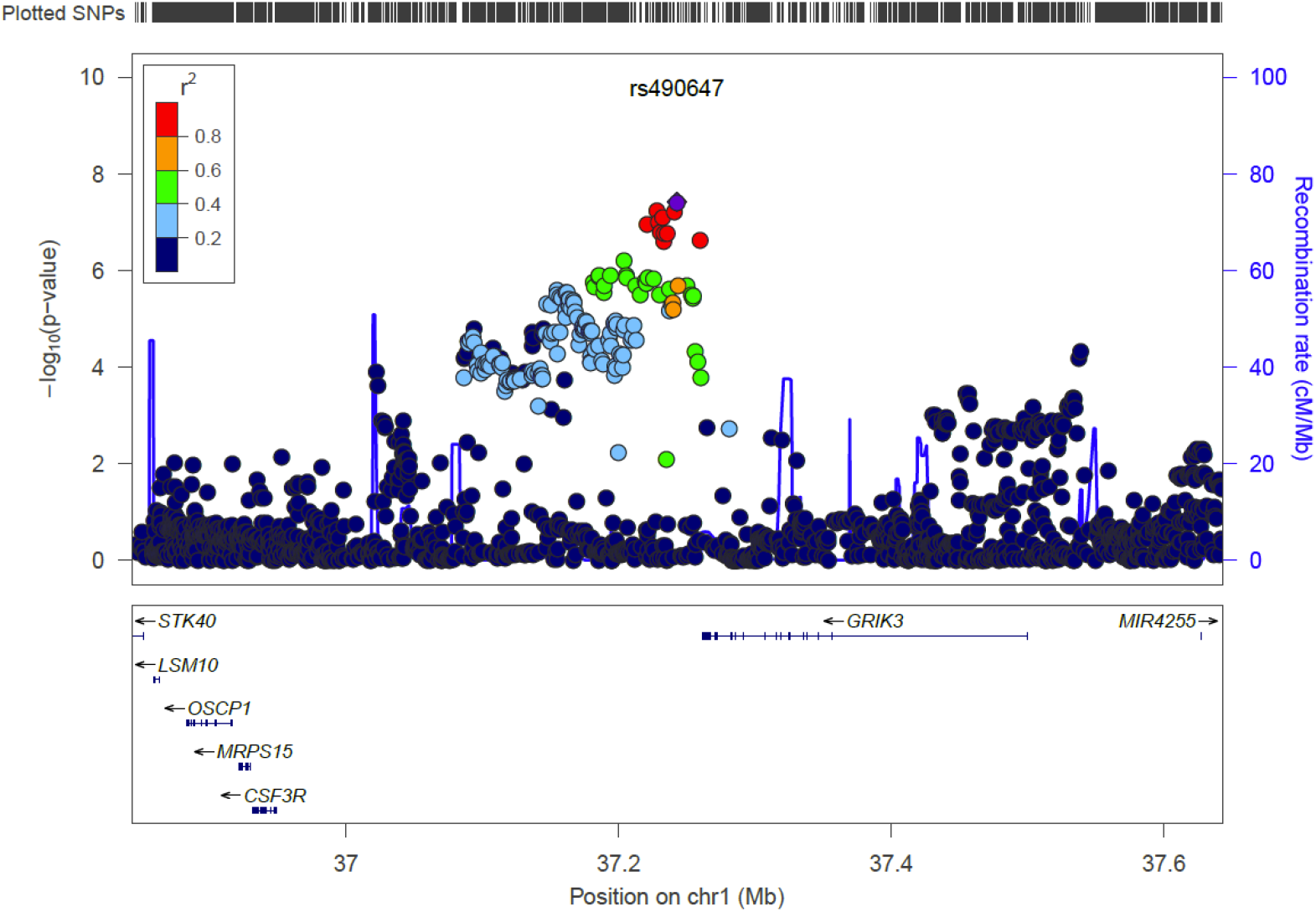
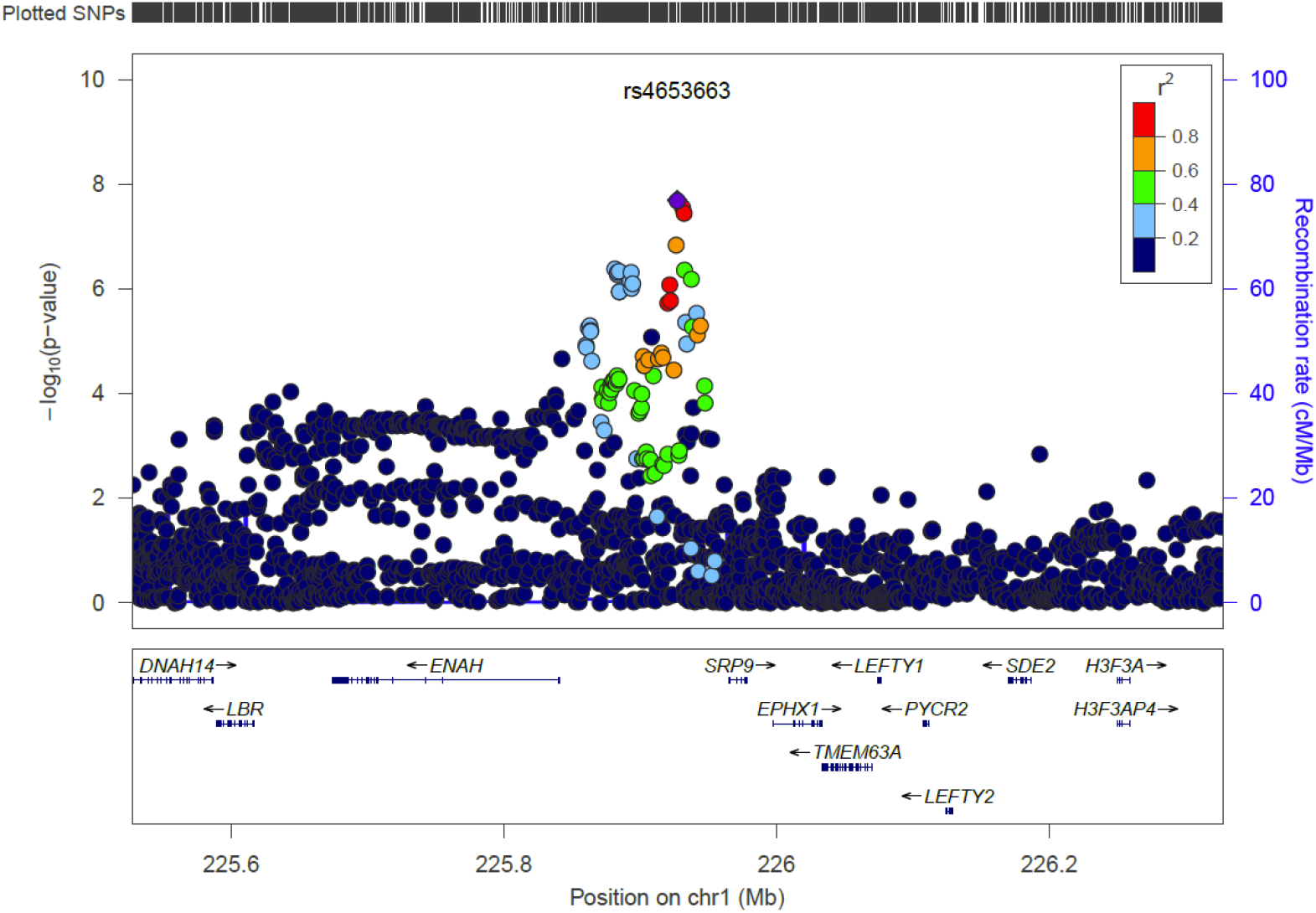
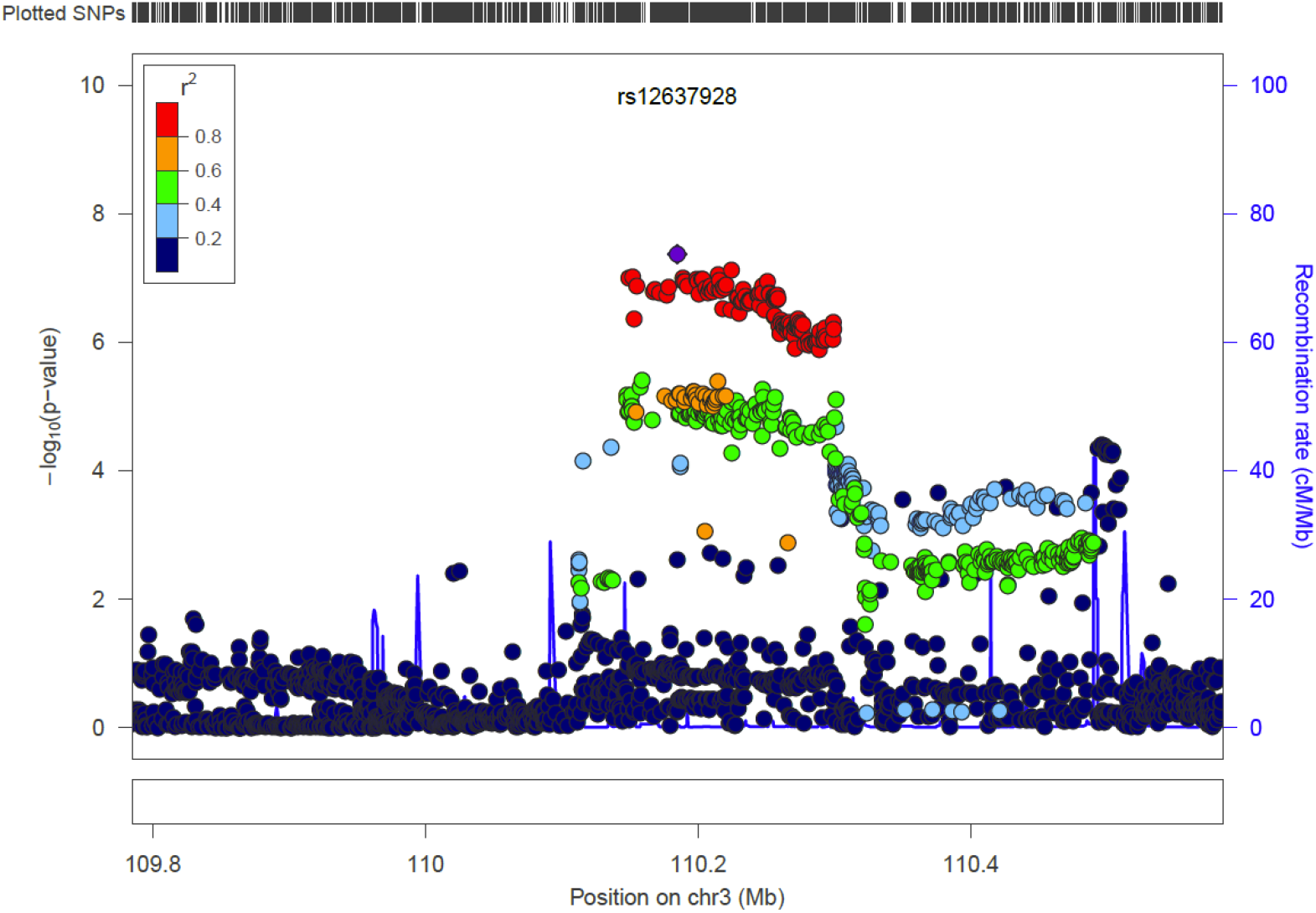
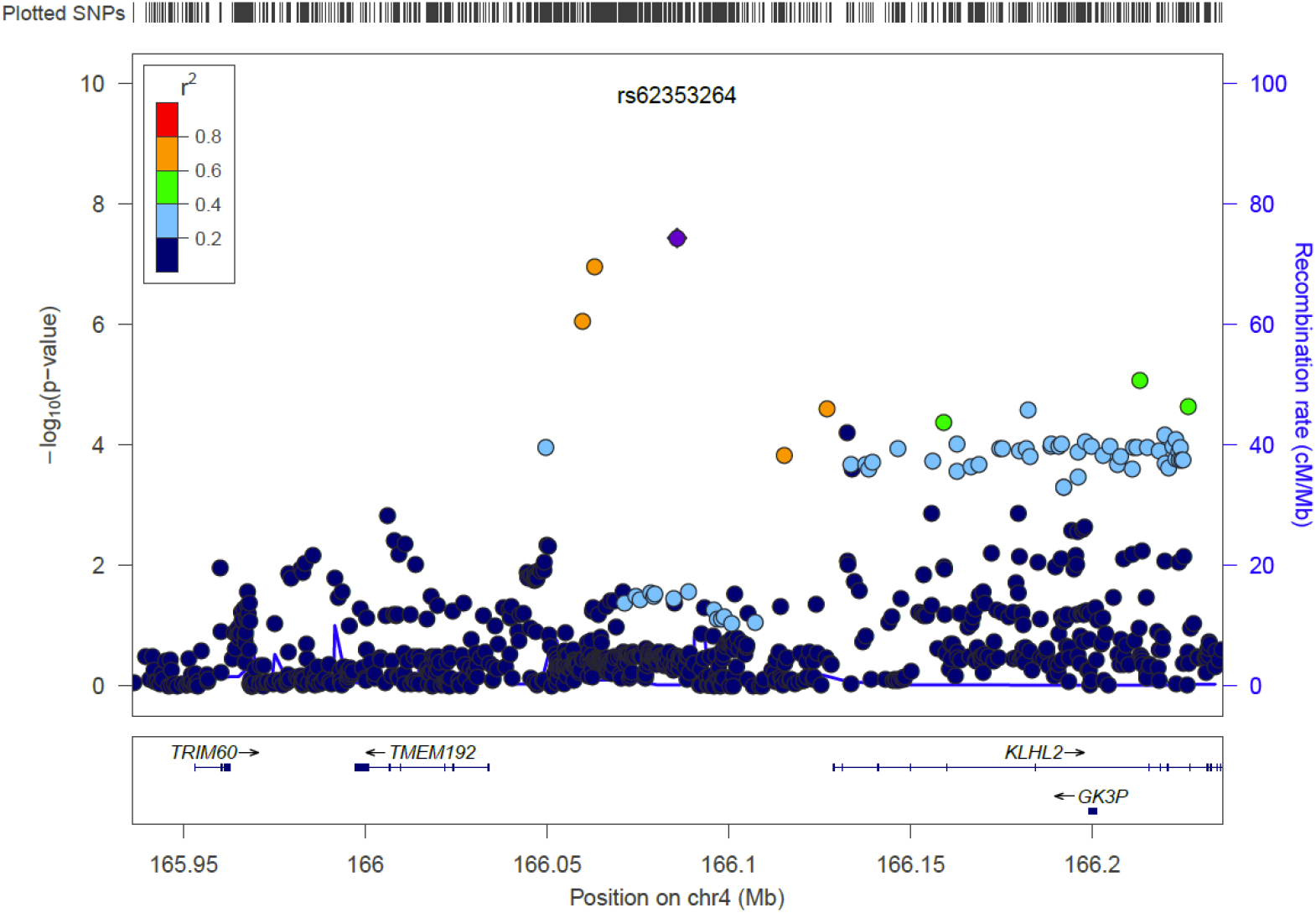
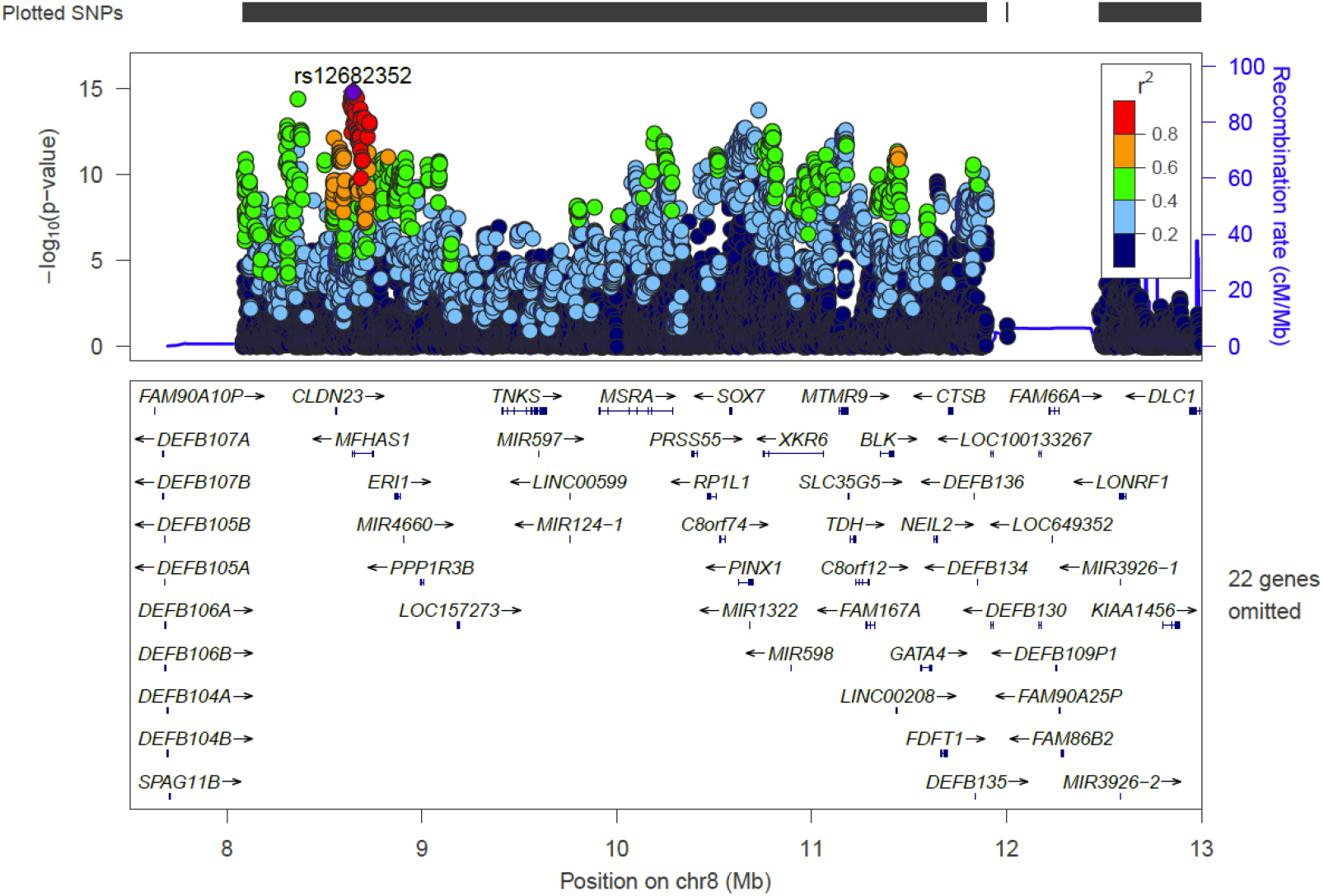
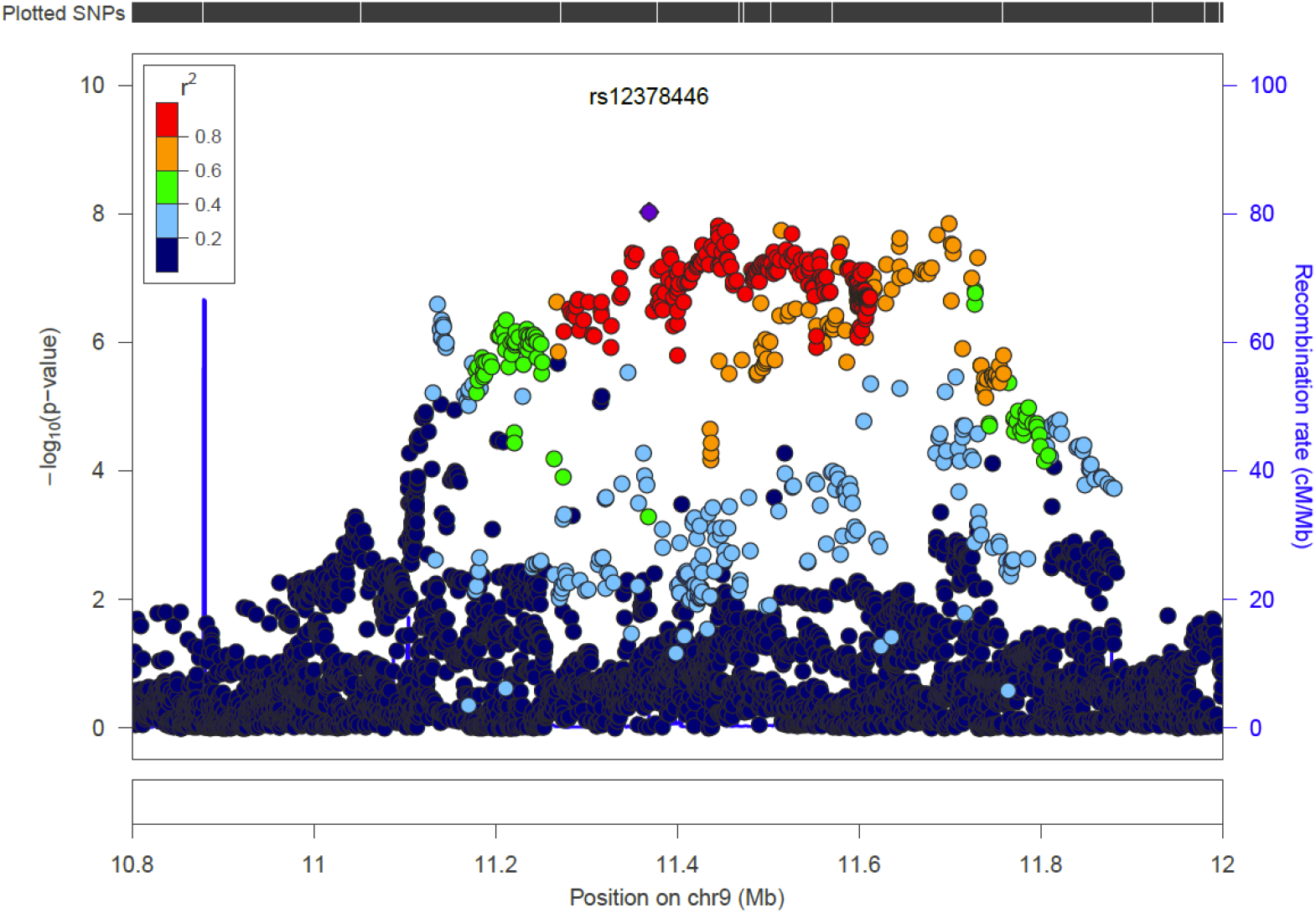
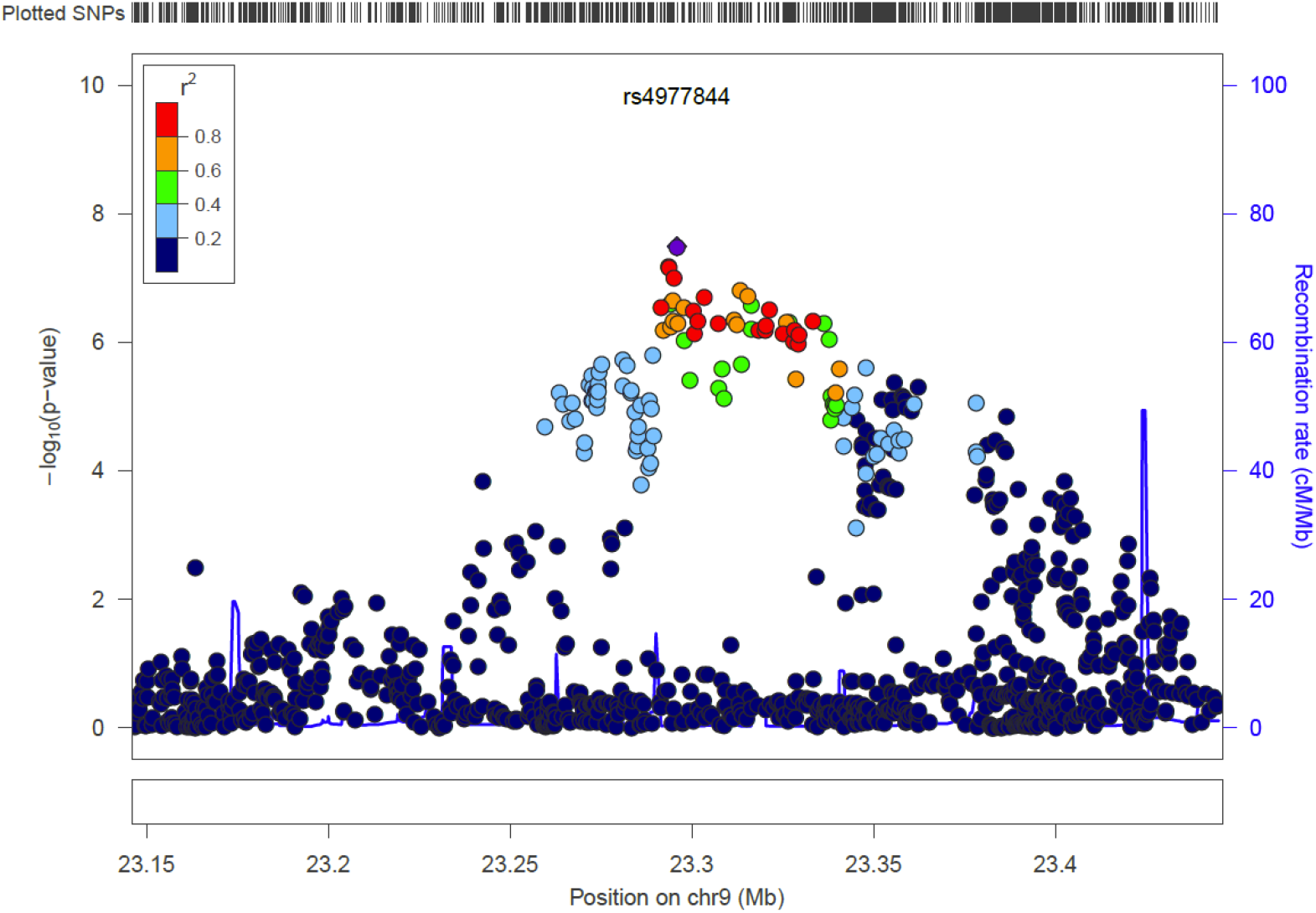
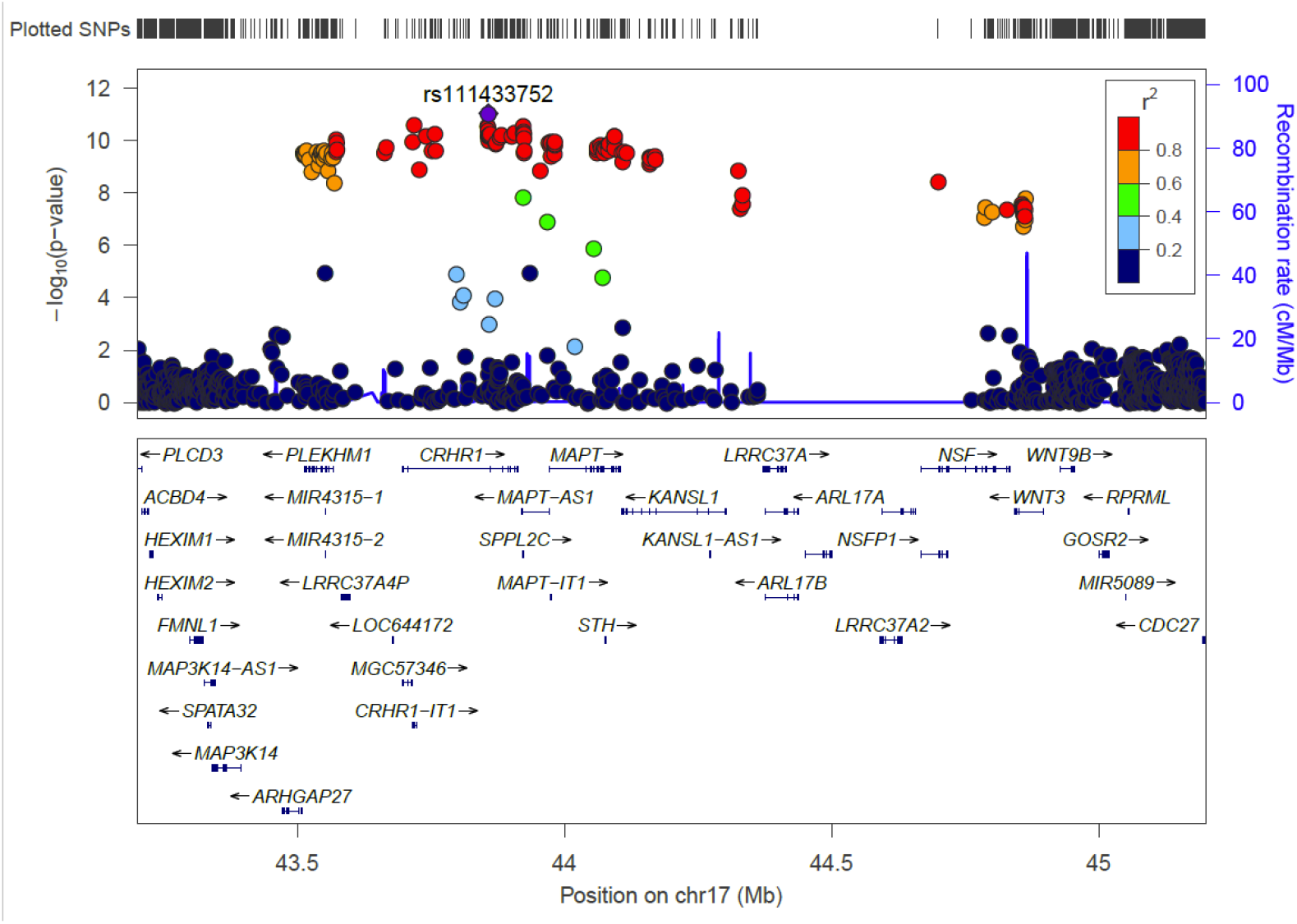
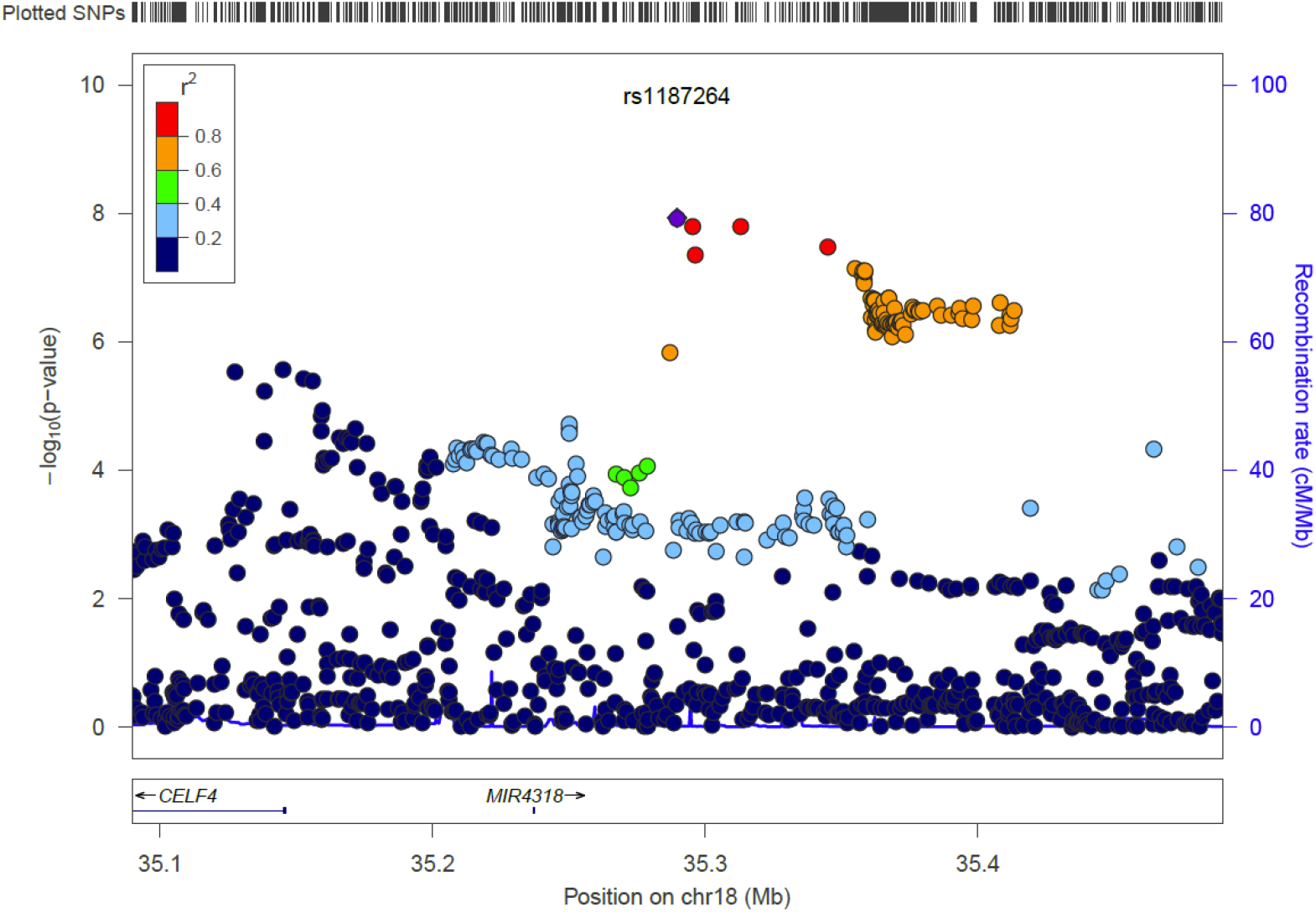
Regional plots of genome-wide significant loci within the meta-analysis of UK Biobank, GS:SFHS and QIMR samples (figures S3a-S3i).

**Figure S4.**
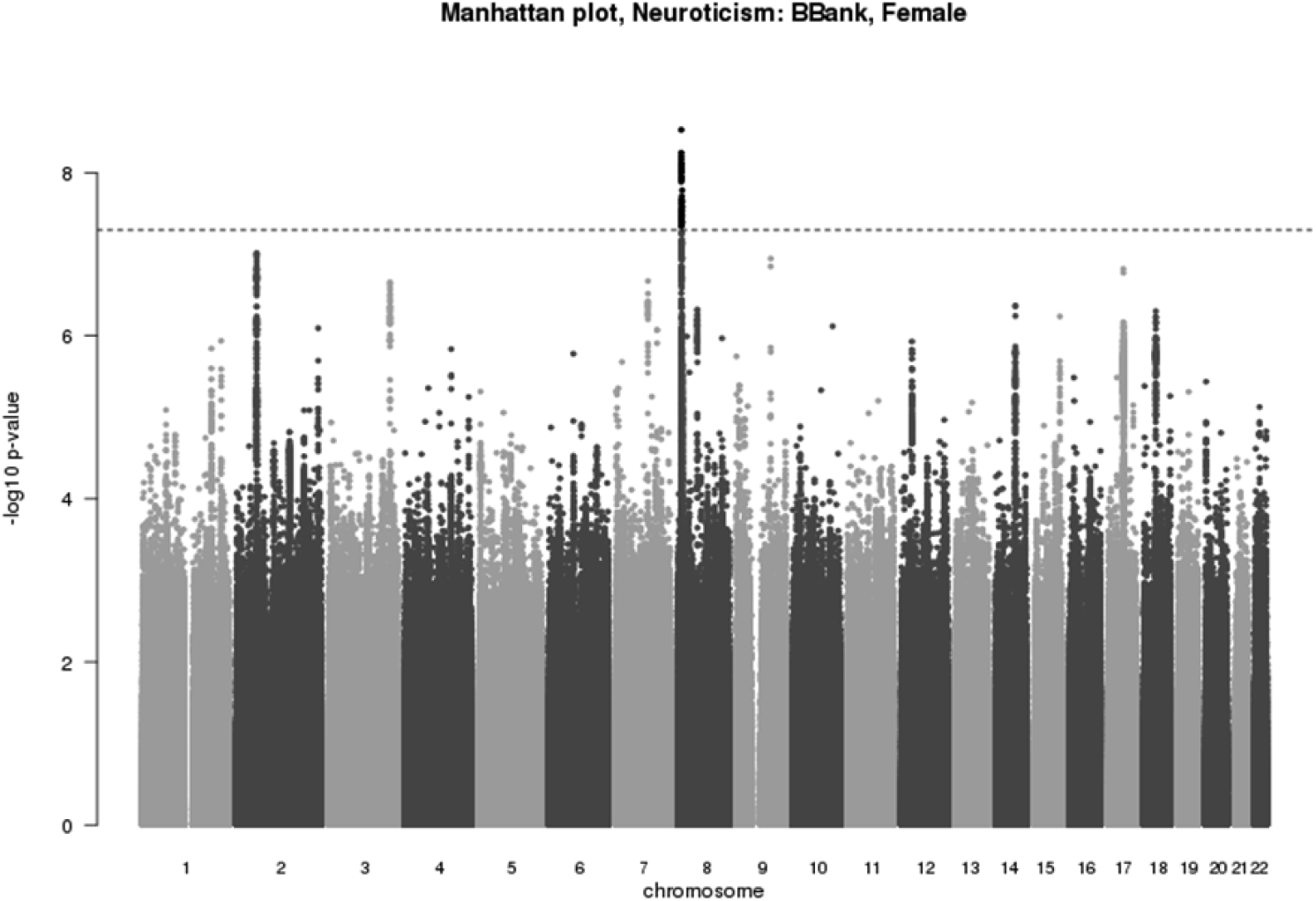
Manhattan plot for genome-wide association with neuroticism in UK Biobank, females only (n=47,196).

**Figure S5.**
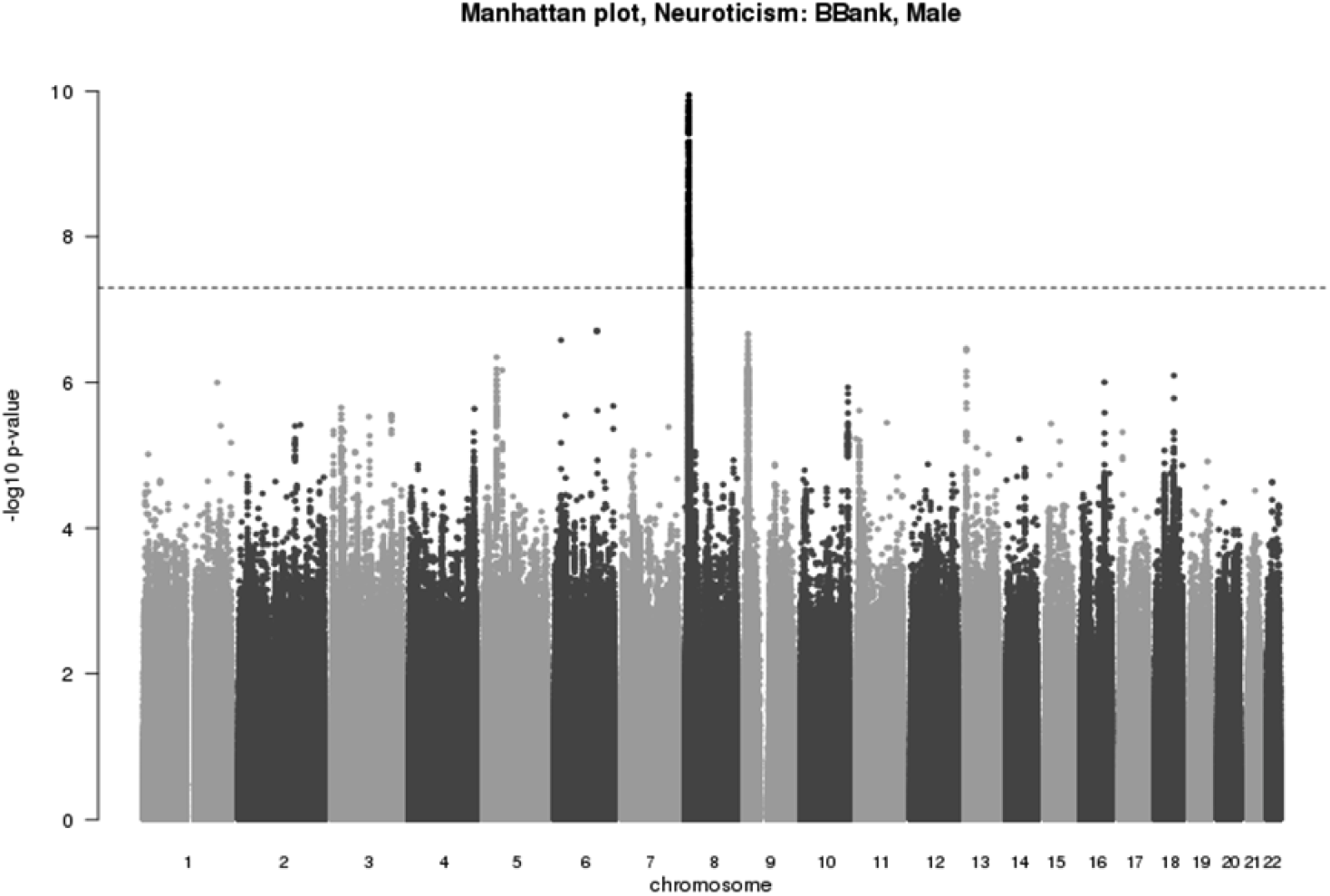
Manhattan plot for genome-wide association with neuroticism in UK Biobank, males only (n=44,174).

**Table S1.** Eysenck Personality Questionnaire-Revised Short Form (EPQ-R-S) Neuroticism scale (24).

|  |  | UK Biobank data-field |
| --- | --- | --- |
| 1 | Does your mood often go up and down? | 1920 |
| 2 | Do you ever feel 'just miserable' for no reason? | 1930 |
| 3 | Are you an irritable person? | 1940 |
| 4 | Are your feelings easily hurt? | 1950 |
| 5 | Do you often feel 'fed-up'? | 1960 |
| 6 | Would you call yourself a nervous person? | 1970 |
| 7 | Are you a worrier? | 1980 |
| 8 | Would you call yourself tense or 'highly strung'? | 1990 |
| 9 | Do you worry too long after an embarrassing experience? | 2000 |
| 10 | Do you suffer from 'nerves'? | 2010 |
| 11 | Do you often feel lonely? | 2020 |
| 12 | Are you often troubled by feelings of guilt? | 2030 |

**Table S2.** Component loadings (on the first unrotated principal component), internal consistency reliabilities and variance explained from principal components analysis of the twelve EPQ-R-S items.

|  |  | **Full UK Biobank**<br>**sample with**<br>**neuroticism data**<br>**(N=401,695)** | **Neuroticism**<br>**GWAS**<br>**sample**<br>**(N=91,370)** |
| --- | --- | --- | --- |
| Item<br>factor<br>loadings | 1. Does your mood often go up and down? | 0.68 | 0.62 |
|  | 2. Do you ever feel 'just miserable' for no reason? | 0.64 | 0.62 |
|  | 3. Are you an irritable person? | 0.52 | 0.64 |
|  | 4. Are your feelings easily hurt? | 0.59 | 0.63 |
|  | 5. Do you often feel 'fed-up'? | 0.66 | 0.62 |
|  | 6. Would you call yourself a nervous person? | 0.61 | 0.63 |
|  | 7. Are you a worrier? | 0.63 | 0.62 |
|  | 8. Would you call yourself tense or 'highly strung'? | 0.60 | 0.64 |
|  | 9. Do you worry too long after an embarrassing experience? | 0.58 | 0.63 |
|  | 10. Do you suffer from 'nerves'? | 0.57 | 0.64 |
|  | 11. Do you often feel lonely? | 0.50 | 0.64 |
|  | 12. Are you often troubled by feelings of guilt? | 0.57 | 0.63 |
| Cronbach's $\alpha$ | | 0.83 | 0.84 |
| % Variance explained by first unrotated principal component |  | 36% | 33% |

**Table S3:** Eight genome-wide significant associations for neuroticism within the UK Biobank dataset.

| Index SNP | Chr | Position | A1/A2 | Frq | BETA (SE) | P | Associated regions |
| --- | --- | --- | --- | --- | --- | --- | --- |
| rs2678897 | 2 | 58,169,418 | G/A | 0.391 | -0.088 (0.016) | $1.45 \times 10^{-8}$ | 57,942,987- 58,484,172 |
| rs62353260 | 4 | 166,078,832 | A/G | 0.013 | 0.361 (0.066) | $3.78 \times 10^{-8}$ | 166,049,663- 166,226,487 |
| rs140344078 | 7 | 7,700,640 | GT/G | 0.172 | -0.113 (0.020) | $1.43 \times 10^{-8}$ | 7,683,347- 7,769,938 |
| rs12682352 | 8 | 8,646,246 | C/T | 0.475 | -0.12 (0.015) | $1.02 \times 10^{-15}$ | 8,088,230- 11,922,801 |
| rs74311404 | 9 | 1,1506,513 | T/TAA | 0.220 | -0.103 (0.018) | $1.58 \times 10^{-8}$ | 11,267,514- 11,810,796 |
| rs8081460 | 17 | 8,965,272 | A/G | 0.307 | -0.091 (0.016) | $2.65 \times 10^{-8}$ | 8,964,004- 8,974,522 |
| rs549599956 | 17 | 44,247,164 | G/A | 0.232 | 0.106 (0.018) | $4.06 \times 10^{-9}$ | 43,501,442- 44,863,133 |
| rs1187256 | 18 | 35,295,330 | T/C | 0.128 | 0.127 (0.023) | $2.16 \times 10^{-8}$ | 35,287,090- 35,413,260 |

*Shown are LD-independent genome-wide significant SNP associations for neuroticism (sorted by genomic position according to UCSC hg19/NCBI Build 37). Column A1/A2 has the SNP alleles, with the first allele ( $\alpha_1$ ) the reference allele for the frequency and BETA columns. Frq=frequency of allele 1. Chr and Position denote the location of the index SNP. BETA=linear regression coefficient for allele1, SE=standard error for BETA. Column 8 notes intersections of all genome-wide significant SNPs positions +/-500KB, with protein-coding genes based on GENCODE gene models (v19, file= gencode.v19.annotation.gtf filtered for feature\_type="gene", gene\_type="protein\_coding" and gene\_status="KNOWN", http://www.gencodegenes.org/releases/19.html).*

**Table S4.**
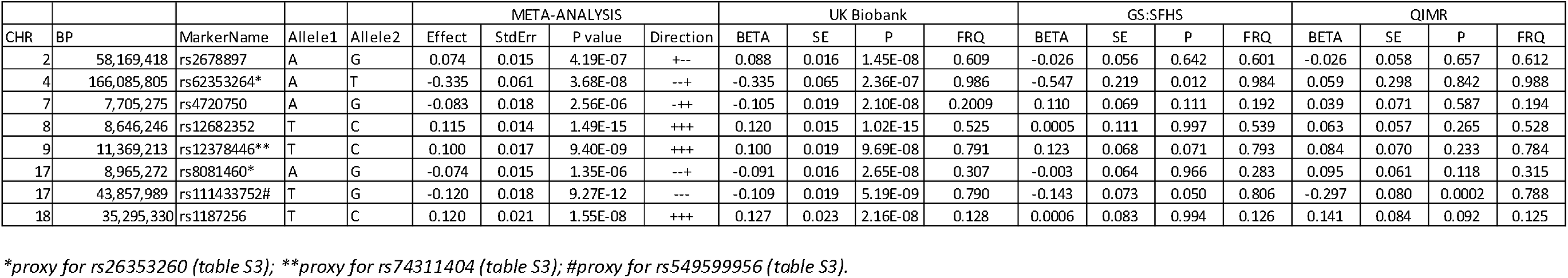
Genome-wide significant index SNPs from UK Biobank (or proxy where not available) within GS:SFHS and QIMR datasets.

## References

1. Matthews G, Deary, I.J., Whiteman., M.C. Personality Traits. 3rd ed. Cambridge. Cambridge University Press. 2009.

2. Wray NR, Birley AJ, Sullivan PF, Visscher PM, Martin NG. Genetic and phenotypic stability of measures of neuroticism over 22 years. Twin Res Hum Genet. 2007;10(5):695-702.

3. Cuijpers P, Smit F, Penninx BH, de Graaf R, ten Have M, Beekman AF. Economic costs of neuroticism: A population-based study. Archives of General Psychiatry. 2010;67(10):1086-93.

4. Weiss A, Gale CR, Batty GD, Deary IJ. Emotionally Stable, Intelligent Men Live Longer: The Vietnam Experience Study Cohort. Psychosomatic Medicine. 2009;71(4):385-94.

5. Kendler KS, Myers J. The genetic and environmental relationship between major depression and the five-factor model of personality. Psychological Medicine. 2010;40(05):801-6.

6. Middeldorp CM, Cath DC, van den Berg M, Beem AL, Van Dyck R, Boomsma DI. The association of personality with anxious and depressive psychopathology. In: Canli T, editor. The Biological Basis of Personality and Individual Differences. New York, NY: Guilford Press; 2006. p. 251-72.

7. Kotov R, Gamez W, Schmidt F, Watson D. Linking “big” personality traits to anxiety, depressive, and substance use disorders: a meta-analysis. Psychological Bulletin. 2010;136(5):768-821.

8. Distel MA, Trull TJ, Willemsen G, Vink JM, Derom CA, Lynskey M, et al. The Five-Factor Model of Personality and Borderline Personality Disorder: A Genetic Analysis of Comorbidity. Biological Psychiatry. 2009;66(12):1131-8.

9. van Os J, Jones, P.B. Neuroticism as a risk factor for schizophrenia. Psychological Medicine 2001;31(06):1129-34.

10. Insel TR, Cuthbert BN. Endophenotypes: Bridging Genomic Complexity and Disorder Heterogeneity. Biological Psychiatry. 2009;66(11):988-9.

11. Cuthbert B, Insel T. Toward the future of psychiatric diagnosis: the seven pillars of RDoC. BMC Medicine. 2013;11(1):126.

12. Eysenck HJ. The biological basis of personality. Springfield, Illinois: Thomas; 1967.

13. Birley AJ, Gillespie N, Heath AC, Sullivan PF, Boomsma DI, Birley AJ, et al. Heritability and nineteen-year stability of long and short EPQ-R Neuroticism scales. Personality and Individual Differences. 2006;40(737-747.).

14. Wray NR, Birley AJ, Sullivan PF, Visscher PF, Martin NG. Genetic and Phenotypic Stability of Measures of Neuroticism Over 22 Years. Twin Research and Human Genetics. 2007;10(5):695-702.

15. Lake RIE, Eave LJ, Maes HHM, Heath AC, Martin NG. Further Evidence Against the Environmental Transmission of Individual Differences in Neuroticism from a Collaborative Study of 45,850 Twins and Relatives on Two Continents. Behavior Genetics. 2000;30(3):223-33.

16. Yamagata S, Suzuki A, Ando J, Ono Y, Kijima N, Yoshimura K, Ostendorf F, Angleitner A, Riemann R, Spinath FM, Livesley WJ, Jang KL. Is the genetic structure of human personality universal? A cross-cultural twin study from North America, Europe, and Asia. J Pers Soc Psychol 90(6):986-98.

17. Keller MC, Coventry WL, Heath AC, Martin NG. Widespread evidence for non-additive genetic variation in Cloninger’s and Eysenck’s personality dimensions using a twin plus sibling design. Behav Genet. 2005;35(6):707-21.

18. van den Berg SM, de Moor MHM, McGue M, Pettersson E, Terracciano A, Verweij KJH, et al. Harmonization of Neuroticism and Extraversion phenotypes across inventories and cohorts in the Genetics of Personality Consortium: an application of Item Response Theory. Behavior Genetics. 2014;44(4):295-313.

19. Genetics of Personality Consortium. Meta-analysis of genome-wide association studies for neuroticism, and the polygenic association with major depressive disorder. JAMA Psychiatry. 2015.

20. Lahey BB. Public health significance of neuroticism. Am Psychol 2009;64(4):241-56.

21. Sudlow C, Gallacher J, Allen N, Beral V, Burton P, Danesh J, et al. UK Biobank: An Open Access Resource for Identifying the Causes of a Wide Range of Complex Diseases of Middle and Old Age. PLoS Med. 2015;12(3):e1001779.

22. Smith B, Campbell H, Blackwood D, Connell J, Connor M, Deary I, et al. Generation Scotland: the Scottish Family Health Study; a new resource for researching genes and heritability. BMC Medical Genetics. 2006;7(1):74.

23. CONVERGE consortium. Sparse whole-genome sequencing identifies two loci for major depressive disorder. Nature. 2015;523(7562):588-91.

24. Eysenck SBG, Eysenck HJ, Barrett P. A revised version of the psychoticism scale. Personality and Individual Differences. 1985;6(1):21-9.

25. Costa PT, Jr., & McCrae, R. R. Revised NEO personality inventory (NEO PI-R) and NEO five-factor inventory (NEO-FFI) professional manual. Odessa, FL: Psychological Assessment Resources.; 1992.

26. Gow AJ, Whiteman MC, Pattie A, Deary IJ. Goldberg’s ‘IPIP’ Big-Five factor markers: Internal consistency and concurrent validation in Scotland. Personality and Individual Differences. 2005;39(2):317-29.

27. UK Biobank. Genotype imputation and genetic association studies of UK Biobank, Interim Data Release. 2015 11 September 2015; http://www.ukbiobank.ac.uk/wpcontent/uploads/2014/04/imputation_documentation_May2015.pdf

28. Delaneau O, Zagury, J.-F. & Marchini, J. Improved whole-chromosome phasing for disease and population genetic studies. Nature Methods. 2013;10:5-6.

29. Howie B, Marchini, J., Stephens, M. Genotype imputation with thousands of genomes. G3 (Bethesda). 2011;1:457-70.

30. Huang J, Howie B, McCarthy S, Memari Y, Walter K, Min JL, et al. Improved imputation of low-frequency and rare variants using the UK10K haplotype reference panel. Nat Commun. 2015;6.

31. UK Biobank. Genotyping of 500,000 UK Biobank participants. Description of sample processing workflow and preparation of DNA for genotyping.2015 11 Sept 2015; http://biobank.ctsu.ox.ac.uk/crystal/refer.cgi?id=155581

32. Yang J, Benyamin B, McEvoy BP, Gordon S, Henders AK, Nyholt DR, et al. Common SNPs explain a large proportion of the heritability for human height. Nat Genet. 2010;42(7):565-9.

33. Bulik-Sullivan BK, Loh P-R, Finucane HK, Ripke S, Yang J, Schizophrenia Working Group of the Psychiatric Genomics C, et al. LD Score regression distinguishes confounding from polygenicity in genome-wide association studies. Nat Genet. 2015;47(3):291-5.

34. Bulik-Sullivan B, Finucane HK, Anttila V, Gusev A, Day FR, Loh P-R, et al. An atlas of genetic correlations across human diseases and traits. Nat Genet. 2015;advance online publication.

35. Psychiatric GWAS Consortium Bipolar Working Group. Large-scale genome-wide association analysis of bipolar disorder identifies a new susceptibility locus near ODZ4. Nat Genet. 2011;43(10):977-83.

36. Schizophrenia Working Group of the Psychiatric Genomics Consortium. Biological insights from 108 schizophrenia-associated genetic loci. Nature. 2014;511(7510):421-7.

37. Major Depressive Disorder Working Group of the Psychiatric Genomics Consortium. A mega-analysis of genome-wide association studies for major depressive disorder. Molecular Psychiatry. 2013;18(4):10.1038/mp.2012.21.

38. Wray NR, Lee SH, Mehta D, Vinkhuyzen AAE, Dudbridge F, Middeldorp CM. Research Review: Polygenic methods and their application to psychiatric traits. Journal of Child Psychology and Psychiatry. 2014;55(10):1068-87.

39. Yang JL, Hong S, Goddard ME, Visscher PM. GCTA: a tool for genome-wide complex trait analysis. Am J Hum Genet. 2011;88:76.

40. Davies G, Armstrong N, Bis JC, Bressler J, Chouraki V, Giddaluru S, et al. Genetic contributions to variation in general cognitive function: a meta-analysis of genome-wide association studies in the CHARGE consortium (N=53 949). Molecular Psychiatry. 2015;20(2):183-92.

41. Benjamini Y, Hochberg Y. Controlling the false discovery rate: a practical and powerful approach to multiple testing. J R Stat Soc Series B (Methodological) 1995:289-300.

42. Kendler KS NM, Kessler RC, Heath AC, Eaves LJ. A longitudinal twin study of personality and major depression in women. Archives of General Psychiatry. 1993;50:853-62.

43. Okbay A, Baselmans BML, De Neve J-E, Turley P, Nivard MG, Fontana MA, et al. Genetic Associations with Subjective Well-Being Also Implicate Depression and Neuroticism. bioRxiv. 2015.

44. Kendler KS, Neale MC, Kessler RC, Heath AC, Eaves LJ. A longitudinal twin study of personality and major depression in women. Archives of General Psychiatry. 1993;50(11):853-62.

45. Kendler KS, Gardner CO. Sex Differences in the Pathways to Major Depression: A Study of Opposite-Sex Twin Pairs. American Journal of Psychiatry. 2014;171(4):426-35.

46. Parker G, Brotchie H. Gender differences in depression. International Review of Psychiatry. 2010;22(5):429-36.

47. Sanacora G, Treccani G, Popoli M. Towards a glutamate hypothesis of depression: An emerging frontier of neuropsychopharmacology for mood disorders. Neuropharmacology. 2012;62(1):63-77.

48. Gray AL, Hyde TM, Deep-Soboslay A, Kleinman JE, Sodhi MS. Sex differences in glutamate receptor gene expression in major depression and suicide. Mol Psychiatry. 2015;20(9):1057-68.

49. Wagnon JL, Briese M, Sun W, Mahaffey CL, Curk T, Rot G, Ule J, Frankel WN. CELF4 regulates translation and local abundance of a vast set of mRNAs, including genes associated with regulation of synaptic function. PLoS Genetics. 2012;8(11):e1003067.

50. Stetler C, Miller G. Depression and hypothalamic-pituitary-adrenal activation: a quantitative summary of four decades of research. Psychosom Med. 2011;73:114-26.

51. Gale C, Hagenaars SP, Davies G, Hill WD, Liewald DC, Cullen B, et al. Pleiotropy between neuroticism and physical and mental health: findings from 108 038 men and women in UK Biobank. BioRxiv. 2015.

52. Weber H, Richter J, Straube B, Lueken U, Domschke K, Schartner C, et al. Allelic variation in CRHR1 predisposes to panic disorder: evidence for biased fear processing. Mol Psychiatry. 2015.

53. Ittner LM KY, Delerue F, Bi M, Gladbach A, van Eersel J, Wölfing H, et al. Dendritic function of tau mediates amyloid-beta toxicity in Alzheimer’s disease mouse models. Cell. 2010;142:387-97.

54. Kimura T, Whitcomb DJ, Jo J, Regan P, Piers T, Heo S, Brown C, Hashikawa T, Murayama M, Seok H, Sotiropoulos I, Kim E, Collingridge GL, Takashima A, Cho K. Microtubule-associated protein tau is essential for long-term depression in the hippocampus. Philos Trans R Soc Lond B Biol Sci 2014;369:2013-144.

55. Cantero JL, Moreno-Lopez B, Portillo F, Rubio A, Hita-Yañez E, Avila J. Role of tau protein on neocortical and hippocampal oscillatory patterns. Hippocampus. 2012:827-34.

56. Wang J, Gao QS, Wang Y, Lafyatis R, Stamm S, Andreadis A. Tau exon 10, whose missplicing causes frontotemporal dementia, is regulated by an intricate interplay of cis elements and trans factors. TJ Neurochem. 2004;88(1078-1090.).

57. Chiesa A, Crisafulli C, Porcelli S, Han C, Patkar AA, Lee SJ. Influence of GRIA1, GRIA2 and GRIA4 polymorphisms on diagnosis and response to treatment in patients with major depressive disorder. Eur Arch Psychiatry Clin Neurosci. 2012;262:305-11.

58. Lee PH, Perlis RH, Jung JY, Byrne EM, Rueckert E, Siburian R. Multi-locus genome-wide association analysis supports the role of glutamatergic synaptic transmission in the etiology of major depressive disorder. Transl Psychiatry. 2012;2:e184.

59. Minelli A, Scassellati C, Bonvicini C, Perez J, Gennarelli M. An association of GRIK3 Ser310Ala functional polymorphism with personality traits. Neuropsychobiology. 2009;59:28-33.

60. Paddock S, Laje G, Charney D, Rush AJ, Wilson AF, Sorant AJ. Association of GRIK4 with outcome of antidepressant treatment in the STAR[ast]D cohort. Am J Psychiatry. 2007;164:1181-8.

61. Schiffer HH, Heinemann SF. Association of the human kainate receptor GluR7 gene (GRIK3) with recurrent major depressive disorder. Am J Med Genet B Neuropsychiatr Genet. 2007;144B:20-6.

62. Tsunoka T, Kishi T, Ikeda M, Kitajima T, Yamanouchi Y, Kinoshita Y. Association analysis of group II metabotropic glutamate receptor genes (GRM2 and GRM3) with mood disorders and fluvoxamine response in a Japanese population. Prog Neuropsychopharmacol Biol Psychiatry. 2009;33:875-9.

63. Tseng LA, Bixby J. Interaction of an intracellular pentraxin with a BTB-Kelch protein is associated with ubiquitylation, aggregation and neuronal apoptosis. Mol Cell Neurosci 2011;47:254-64.

64. Soltysik-Espanola M, Rogers RA, Jiang S, Kim TA, Gaedigk R, White RA, Avraham H, Avraham S.. Characterization of Mayven, a novel actin-binding protein predominantly expressed in brain. Mol Cell Biol 1999;10:2361-75.

65. Jiang S, Avraham HK, Park SY, Kim TA, Bu X, Seng S, Avraham S. Process elongation of oligodendrocytes is promoted by the Kelch-related actin-binding protein Mayven. J Neurochem 2005;92:1191-203.

66. Takahashi H, Craig AM. Protein tyrosine phosphatases PTPδ, PTPσ, and LAR: presynaptic hubs for synapse organization. Trends Neurosci 2013;36:522-34.

67. Schormair B, Kemlik D, Roeske D, Eckstein G, Xiong L, Lichtner P, et al. PTPRD (protein tyrosine phosphatase receptor type delta) is associated with restless legs syndrome. Nat Genet. 2008;40:946-8.

68. Verweij KJ, Yang J, Lahti J, Veijola J, Hintsanen M, Pulkki-Råback L. Maintenance of genetic variation in human personality: testing evolutionary models by estimating heritability due to common causal variants and investigating the effect of distant inbreeding. Evolution. 2012;66(10):3238-51.

69. Vinkhuyzen AAE, Pedersen NL, Yang J, Lee SH, Magnusson PKE, Iacono WG, et al. Common SNPs explain some of the variation in the personality dimensions of neuroticism and extraversion. Transl Psychiatry. 2012;2:e102.

70. Barlow DH, Ellard KK, Sauer-Zavala S, Bullis JR, Carl JR. The Origins of Neuroticism. Perspectives on Psychological Science. 2014;9(5):481-96.

71. Van Os J, Park, S.B.G., & Jones, P.B. Neuroticism, life events and mental health: evidence for person-environment correlation. British Journal of Psychiatry. 2001;178 (suppl. 40):s72-s5.

72. Insel T. The NIMH Research Domain Criteria (RDoC) Project: Precision Medicine for Psychiatry. American Journal of Psychiatry. 2014;171(4):395-7.

